# Mitochondrial Genome Diversity in the Central Siberian Plateau with Particular Reference to Prehistory of Northernmost Eurasia

**DOI:** 10.1101/656181

**Authors:** S. V. Dryomov, A. M. Nazhmidenova, E. B. Starikovskaya, S. A. Shalaurova, N. Rohland, S. Mallick, R. Bernardos, A. P. Derevianko, D. Reich, R. Sukernik

## Abstract

The Central Siberian Plateau was last geographic area in Eurasia to become habitable by modern humans after the Last Glacial Maximum (LGM). Through comprehensive mitochondrial DNA genomes retained in indigenous Siberian populations, the Ket, Tofalar, and Todzhi - we explored genetic links between the Yenisei-Sayan region and Northeast Eurasia over the last 10,000 years. Accordingly, we generated 218 new complete mtDNA sequences and placed them into compound phylogenies along with 7 newly obtained and 70 published ancient mt genomes. Our findings reflect the origins and expansion history of mtDNA lineages that evolved in South-Central Siberia, as well as multiple phases of connections between this region and distant parts of Eurasia. Our result illustrates the importance of jointly sampling modern and prehistoric specimens to fully measure the past genetic diversity and to reconstruct the process of peopling of the high latitudes of the Siberian subcontinent.

---

Although modern humans began colonizing sub-arctic and arctic Siberia 45,000 years ago, most archaeological sites postdate 12 kya, with the exception of episodic incursions to the north of Eurasia during the warm phases, with the settlements primary limited to cryptic refugia both in eastern Europe and Asia (reviewed by Pitulko et al., 2017 [1]). During the last millennium, the Central Siberian Plateau, bounded by the Yenisei River to the west, the North Siberian Lowland to the north, the Verkhoynsk Mountains to the east, and the Sayan Mountains and Lake Baikal to the south, has been shaped by key episodes that have had major impacts on human migrations, admixture, and population replacement. At the end of the 15^th^ century, when Russians first crossed the Ural Mountains in substantial numbers, dozens of indigenous tribes and bands with markedly different ways of life and subsistence were widespread throughout Siberia [2,3]. Soon after, some of these groups became extinct or underwent marked postcontact decimation and admixture while others only diminished in number, their territories shrinking to small patches of their former distributions. Two strata are recognized in the aboriginal Siberian population: an ancient layer and a comparatively new one, or the “Old Siberians” versus “Neosiberians” [4,5]. Most of the “Old Siberians” who spoke languages belonging to linguistic outliers, such as Omok (an extinct Yukaghir language spoken as late as the 18^th^ century in the lower Kolyma and Indigirka valleys), have been greatly reduced in number, and their cultures have been brought under the domination of the more populous Neosiberians: Tungusic, Turkic or Mongolian tribes, all relative newcomers to the region [2,3,5,6]. Nonetheless, groups of Old Siberians still persist in remote “pockets” of Siberia in the form of anthropological isolates, usually representing the now-amalgamating remnants of small, previously distinct tribes and lineages. The study of mitochondrial DNA variation to ascertain their origin and affinities, as well as their relationship to the first Americans, has been a subject of intensive study since the early 1990s [7-9]. Candidate founding lineages for Native American mtDNA haplogroups were identified by identifying similar haplogroups in Eurasia [10-14]. For instance, unique mtDNA sequences (haplogroups B4b1a, D4h2, and C1a) linked to Native American B2, D4h3a, and C1, respectively, were attested using complete mtDNA sequences [10-12,14]. A complete catalogue of mtDNA variation in the “Old-Siberians” is therefore important not just for understanding the specific history of Siberia, but also the ancestry of peoples of the New World.

A central concern of this study is disentangling ancestries from different sources that mixed at different time points in the past to provide a better understanding of the genetic interactions between and within Siberian populations. Accordingly, we focused on mitochondrial DNA genome diversity in the Yenisei-Sayan autochthonous populations, primarily the Ket, Tofalar, and Todzhi (Fig. 1). Ket are now the sole surviving member of the Yeniseian language family, speaking a unique language that does not easily fit into any known phyla and is unrelated to any other Siberian language [15]. Their related tribes, the Assan, Arin, and Kott, disappeared about 200 years after their first contact with Russians. In this region, the Ket is the last tribe to retain their original language and until recent times subsisted entirely on hunting, fishing, and the gathering of wild plants [2,16,17]. The Tofalar and Todzhi languages whose members originally spoke Samoyed belong to the subgroup of Turkic languages confined to the upper Yenisei area [18-20], where they may have had ample opportunity to exchange genes with the Ket or related tribes (Lopatin, 1940 [5], and references therein). We compared mitochondrial DNA present-day diversity from these groups with complete mitochondrial genomes from ancient samples from the region and placed them into combined genealogical trees. We used these updated genealogies to trace ancestral relationships between populations sharing subhaplogroup-specific mutations. We used the results to reconstruct expansions (preponderantly from south to north) as well as contractions of populations, thus shedding new light on northeastern Eurasia’s past.

**Figure 1.**
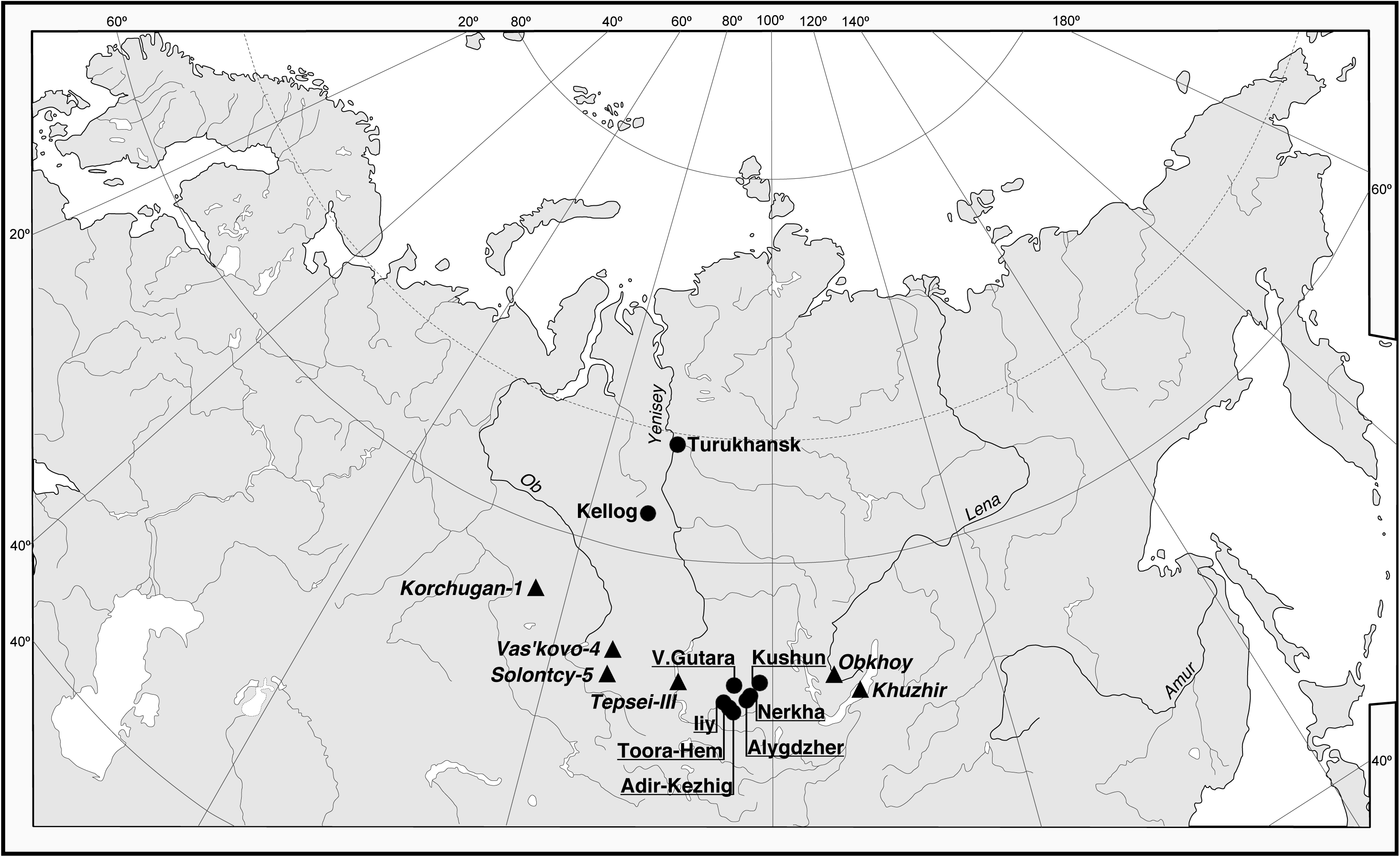
Map of Siberia and adjacent part of Europe. Black circles mark the settlements of sampling expeditions. Black triangles denote locations of the ancient specimens generated through the course of this study and listed in Supplementary Table 2.

## Populations and samples

*Ket -* Formerly called Yenisei Ostyak (the census of 1897 recorded 988 persons), the Ket people are thought to be descendants of some of the earliest inhabitants of Central Siberia, while all of their present-day neighbors seem to be relative newcomers. In the 18^th^ century part of the Ket were forcibly moved to the lands between the Ob and Yenisei Rivers, where the Selkup belonging to the Samoyed group of the Uralic language family lived [2,5,16]. Until recently, there were ∼500 Ket living in a few riverside villages in the middle reaches of the Yenisey; as in the past, many survive as seasonal hunters, trappers, and fishermen. The remnants of the Southern (Upper) Ket, whose ancestors are believed to have originated from the Podkamennaya Tunguska region, have been almost completely integrated into the expanding Evenki [17].

We integrated our newly obtained mtDNA sequences from previously collected DNA samples with previously published data [21] and collected new DNA samples from indivduals residing in August of 2014 in the villages of Turukhansk and Kellog (Turukhansk District, Krasnoyarsk Region, Russian Federation). The total Ket sample subjected to complete mtDNA sequencing was 38 individuals. Based on self-reported ancestry, verified by senior members of related families, we determined that the overwhelming majority of the Ket have been extensively admixed with Russians. A substantial proportion of the Ket were not completely certain of their maternal ancestry in terms of Ket, Selkup, or Evenki origin.

### Tofalar

Fewer than 400 Tofalar, who are reindeer breeders and hunters, inhabit the northern slopes of the eastern Sayan Mountains, along the rivers Uda, Gutara, and Nerkha. They originally spoke a Samoyed language, but later changed to a language of the Turkic family. The Tofalar are comprised of the remnants of several hunting tribes, gradually assimilated into a broader group. In the 17^th^ century, the predecessors of the Tofalar entered into five administrative settlements of “Udinsk Land” of Krasnoyarsk Region [2]. When the Tofalar transitioned to a settled lifestyle in 1930, they were concentrated in three newly established villages in the territory of Nizhni Udinsk District encompassing Upper Gutara, Nerkha, and Alygdzher. There is anthropological and linguistic evidence that classifies Tofalar among the ‘OldSiberians’ [5]. In this study, we revised the analysis of previously collected Tofalar mtDNAs from Upper Gutara and Nerkha villages [10] and supplemented existing data with new blood samples from the easterenmost village of Alygdzer collected in July 2015. The total Tofalar/Udinsk Buryat sample subjected to complete mtDNA sequencing constituted 40 individuals.

### Todzhi

This is a small subgroup living in Todzhinsky District in the northeast part of the Tuva Republic encompassing the intermountain Todzhinsky Depression between the Western and Eastern Sayan. Although modern Todzhi people speak a dialect of the Tuvan language, anthropologically they are distinct from southern and western Tuvans, being similar to the neighboring Tofalar people [18]. Census records of 2002 noted about 3000 Todzhi, and their traditional economy is generally considered to have had a hunting-gathering emphasis supplemented by reindeer herding and fishing. In this study, we focused on 160 blood samples collected through fieldwork in February 2017 in three main settlements of the Todzhinsky District (Toor-Khem, Adyr-Kezhig, and Iy). We selected unrelated samples from elders (n=51) born in 1962 and/or earlier for full mtDNA sequencing.

## RESULTS

In what follows we describe the genetic diversity of complete mtDNA sequences from the Ket, Tofalar, and Todzhi, and compare them with previously reported data. We also combined this with mtDNAs from the Mansi, Tubalar, Nganasan, Evenki, Even, Yukaghir, and Koryak, many of whom we sequenced to the full mtDNA genome level, building on lesser amounts of data previously reported from these samples. A brief description of each population, as well as details of the sampling collection are reported in the work of Torroni et al. 1993b [8]; Derbeneva et al. 2002a, b [21,22]; Starikovskaya et al. 2005 [10]; Volodko et al. 2008 [11]; Sukernik et al. 2010, 2012 [12,23].

### Ket in their Eurasian context

#### Haplogroup N2a

The Yenisei region was outside the main routes of Eurasian agricultural exchange up to the time of the Late Bronze Age Karasuk Culture [24-26]. One of the remarkable features of the Ket mtDNA pool is a lineage of hapogroup N2a distinguished by a set of mutations that we newly document here (m.1633T>C, m.11722C>T, and m.12192G>A) (Fig. 2). Whereas the previously published N2a sample uniquely marked by m.10841A>G (EU787451) is from the Upper Ket in the village of Sulamai on the Stony Tunguska River [21], our newly identified N2a mitogenome lacks this mutation, and comes instead from the village of Kellog in the Lower Ket. Apart from the Ket individuals, none of the other extant or ancient Siberian populations sampled to date are known to carry N2a. This haplogroup is found in just a few contemporary individuals from Europe, Iran, Arabia, and Ethiopia [27,28]. It is likely that this lineage came to the mid-Yenisei from Caucasus, presumably the major source of N2a, which contributed to the central Siberian maternal lineages.

**Figure 2.**
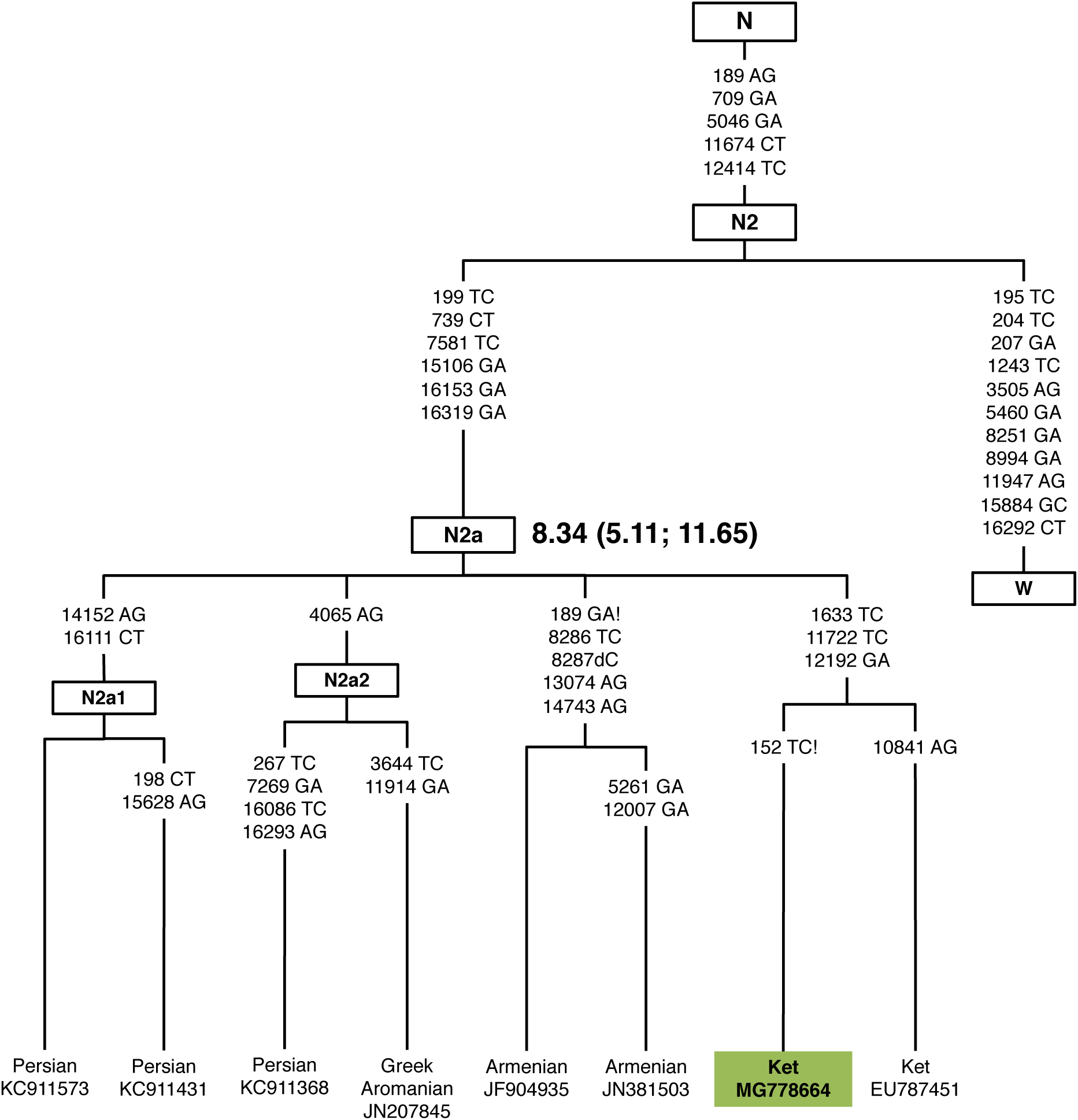
The phylogeny of haplogroup N2a complete sequences.

#### Haplogroup U4a/U4a1

Haplogroup U mtDNAs, confined to subhaplogroups U2e1, U3, U4a, U4b, U4c, U4d, U5a1, is well represented in Old Siberian groups in the northern Altai-lower Ob-Yensei triangle [11,12,21,22,29] including in our newly reported data. We built a phylogenetic tree selectively for U4a/U4a1 subhaplogroups, including ancient mtDNA sequences and their Ket counterparts, or those from the regions close to the Ket traditional territory (Fig. 3). The entire tree, combining the Ket, Tubalar, Mansi, and Vadei Nganasan sequences with ancient mt genomes, a newly reported genome from Novosibirsk (I0992: 5002-4730 calBCE), and a newly reported genome from the neighboring Kemerovo region (I2074:5602-5376 calBCE), allows us to suggest that the U4a1 lineage was present by Neolithic times in Altai-Sayan, where it could have diversified in situ, with U4a1 and U4a3 dispersing into the heartland of Europe. Complete U4a1 mtDNA sequence in the Yamnaya (I0231) [30] culture (the Beaker people of western Europe in the early Bronze Age represented the far western extent of Yamnaya ancestry spread) from Samara region and Mesolithic genomes from Sweden is consistent with the association of the U4a lineage in West Siberia with Mesolithic ancestry.

**Figure 3.**
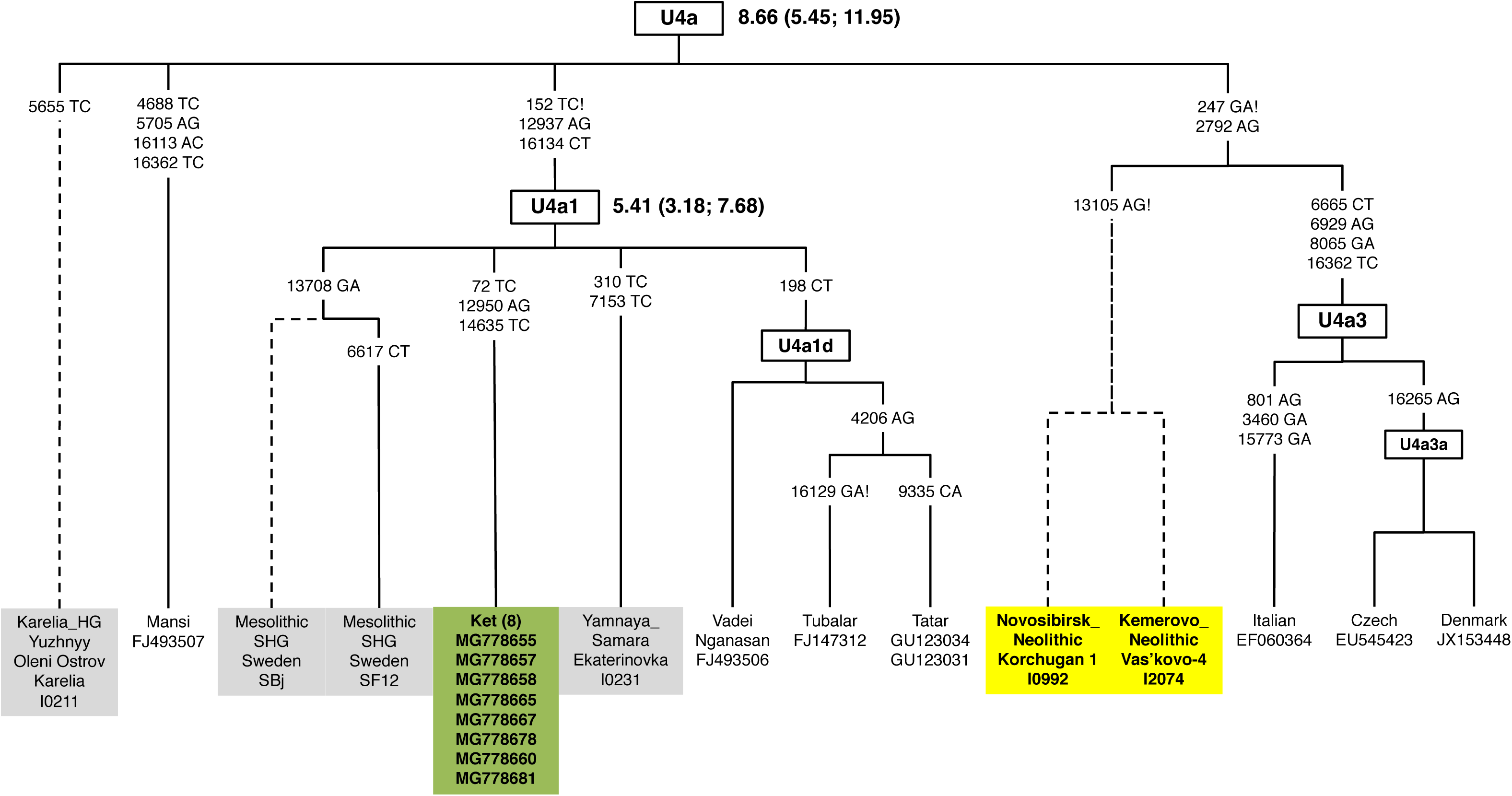
The phylogeny of U4a distinct branches and subbranches.

#### Haplogroup U4d

The updated phylogeny of U4d includes both its main subhaplogroups U4d1 and U4d2, and three previously unreported sublineages documented in two Ket and two Mansi mt genomes (Fig. 4). Whereas the U4d1 sequences are prevalent in the Baltic Sea region, the U4d2 haplogroup is associated with various tribes in areas where eastern Uralic-speakers [11] expanded. Specifically, the observation of U4d1 sequence in a Bronze Age individual (RISE500) from Kytmanovo (along the Chumysh River in the northwestern Altai) supports the hypothesis of an east-to-west pattern of divergence of Uralic languages. The coalescent time of the U4d1 and U4d2 subclusters (∼4.7 kya) is consistent with the hypothesis of continuous gene flow between the Yenisei River valley and the eastern Baltic region during and following the Bronze Age. This conclusion is in agreement with ancient genome-wide data from Finland and the Russian Kola Peninsula [31], revealing that the specific genetic makeup of northern Europe traces important ancestry from Siberia migration that began at least 3,500 years ago.

**Figure 4.**
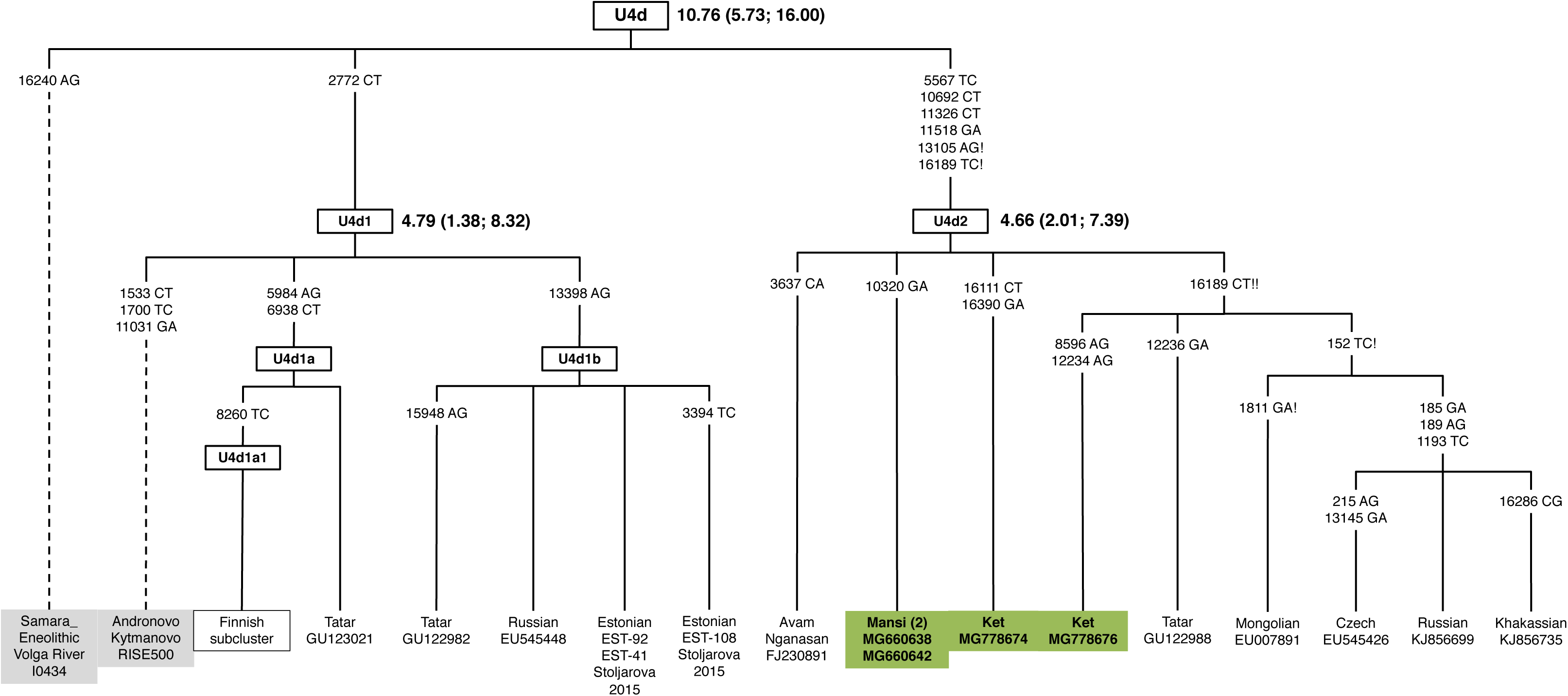
The phylogeny of U4d complete sequences; compared with a couple of ancient (incomplete) sequences.

#### Haplogroup U5a

Recent studies utilizing the genome-wide approach suggest that U5a lineages already existed in Mesolithic Fennoscandia (U5a1 and U5a2) and may imply an eastern origin for these sublineages in Europe [32,33]. Three of our new Ket sequences clustered into U5a1d2 along with a newly reported ancient mt genome (I2068: 420-565 calCE) from the Yenisei-Sayan area, whereas two of the Tubalar sequences fall into U5a1d2b. Previously unrecognized haplogroup U5a2 haplotypes (U5a2b and U5a2a1b) were identified in a few Mansi mtDNAs, chosen from previously collected samples [22]. With a novel U5a1h harbored by a Mansi individual, these findings revise our understanding of U5a1 haplogroup relationships (Supplementary Table 1, Fig. 5).

**Figure 5.**
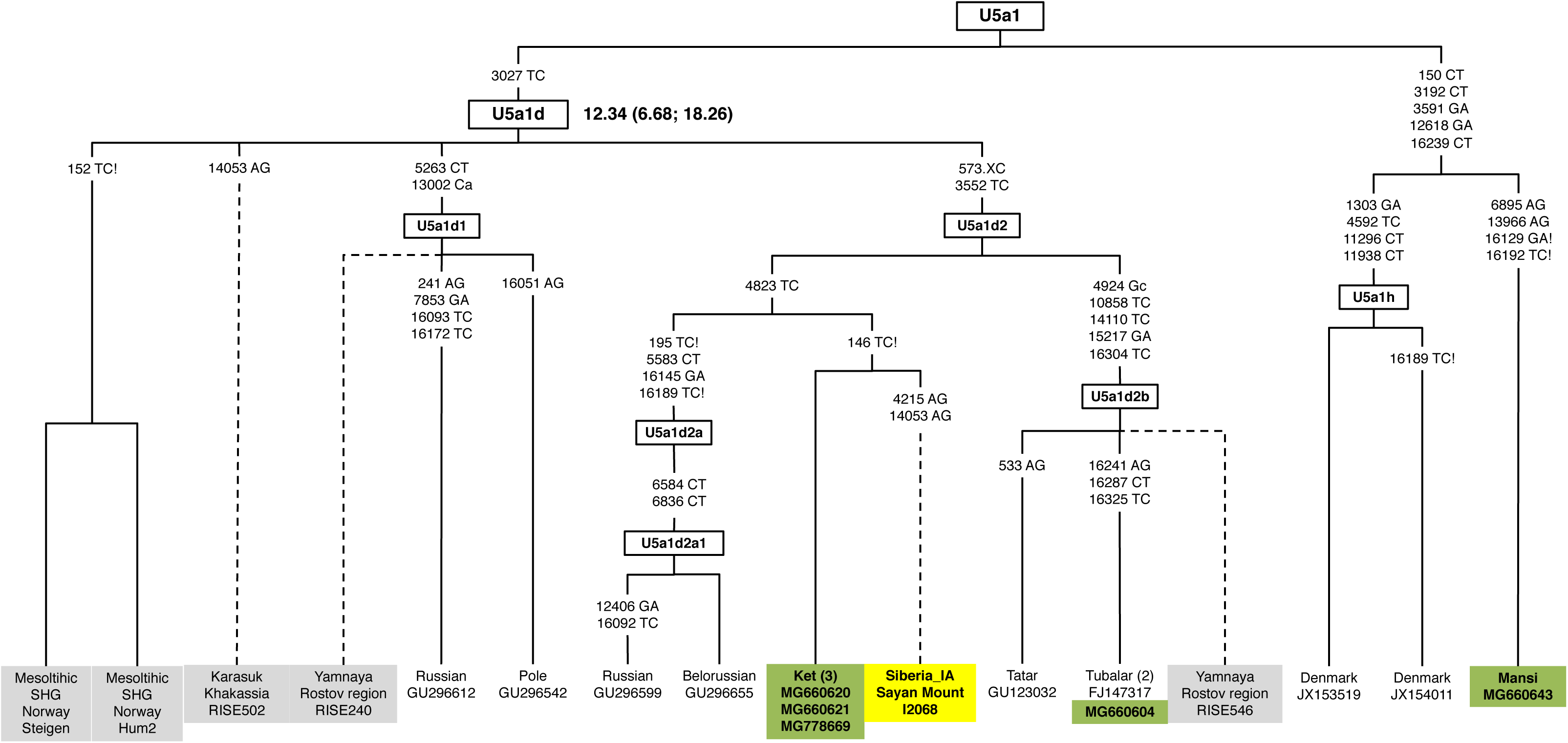
The phylogeny of U5a1 revealing distinct branches or subbranches in the Ket, Tubalar, and Mansi mt genomes compared with ancient sequences.

#### Haplogroup A8

Through the course of this study, previously published and newly obtained A8a2 samples were explored to redefine the structure of the entire A8 tree (Supplementary Table 1; Fig. 6). As a result, we were able to delineate a previously uncategorized A8b lineage [34]. The immediate split of A8 created a distinctive A8b evident in two Koryak mt genomes. The entire tree expands our understanding of haplogroup A8 by encompassing previously attested ancient mt genomes attributed to the Bronze Age Okunevo culture in Khakassia. Hence, genetic and archeological evidence support a single/common origin of a population directly related to the maternal ancestors of some of the present-day Ket, Tofalar, Tuvan, Yakut, Buryat, as well as the Koryak individuals. It has been suggested that the Okunevo Culture derived ancestry from long-resident populations of the Altai-Sayan Upland whose roots may extend back to the Neolithic, if not before [35].

**Figure 6.**
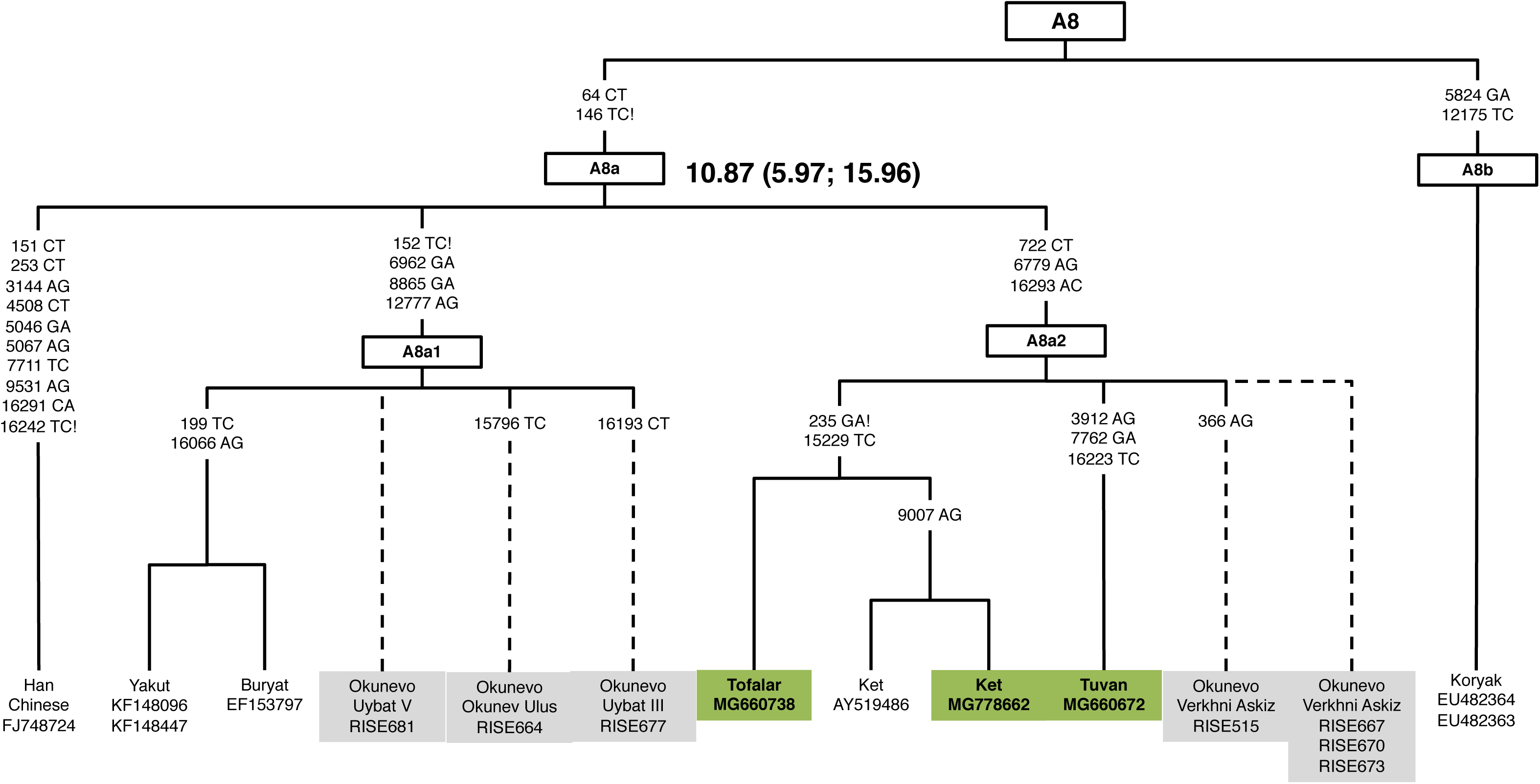
The phylogeny of A8 revealing distinct lineages or sublineages.

### Mitochondrial DNA gene pool in Tofalar and Todzhi

The present-day Tofalar and Todzhi do not cluster with the Russian Buryat who live just to the northeast of Mongolia. Unlike the Ket, the Tofalar and Todzhi showed high frequencies (>50%) of the east Eurasian haplocluster C4a’b, a pattern observed in our previous studies of the Tofalar and Evenki [10]. In most cases, the results initially obtained by examination of a part of the C4a and C4b sequences have been confirmed and extended by subsequent surveys based on complete mitochondrial DNA analysis [11]. Furthermore, eight of the Tofalar/Udinsk Buryat sample (20%) harbored mitogenomes assigned to haplogroup M8a. This lineage, as well as its main derivatives (M8a1, M8a2, M8a3), has not been observed among circumpolar groups, being confined to southern and southeastern Siberia (Supplementary Fig. 1). The geographic origin of M8 root appears somewhere within South-East Asia, subsequently splitting into M8a and CZ clusters in the area currently inhabited by modern East Asians [36].

#### Haplogroup C4

The age estimates of the newly described sublineages hint to the origin and diversification of the C4 haplogroup in Siberia/Asia (Fig. 7). Thus, modern samples (C4d and C4e) nested in C4-m.152T>C node together with the newly reported ancient individual I2072 (Solontsy5 from Altai dating to 3959-3715 calBCE) and RISE602 samples (Sary-Bel, 9/700 BC – AD 500/1000). Likewise, I1000 from Obkhoy, from the Glazkovskaya culture and dated to 2871-2497 calBCE, all clustering to C4, support ancient gene flow within southern extent of the Central-Eastern Siberia. The coalescence time estimates of the C4-m.152T>C-m.16093T>C were estimated to be around 20 kya. Unlike the C4e subhaplogroup revealed in northwestern Altai, the C4d lineage found outside Siberia shared common ancestry with the Tibetans or populations currently living with Tibetans [37,38]. On the other hand, C4c, a sister group of C4a and C4b [11] is found solely in the Western Hemisphere. The C4c present-day distribution in North America is centered on the southern end of the former glacial ice-free corridor. If this is a significant signature of its early dispersal history, it could plausibly have been carried by the earliest peoples of central North America, who are argued on genetic [39] as well as archaeological and paleoecological grounds to have expanded southward from Beringia/Siberia into Americas sometime before 13,500 years ago [40,41].

**Figure 7.**
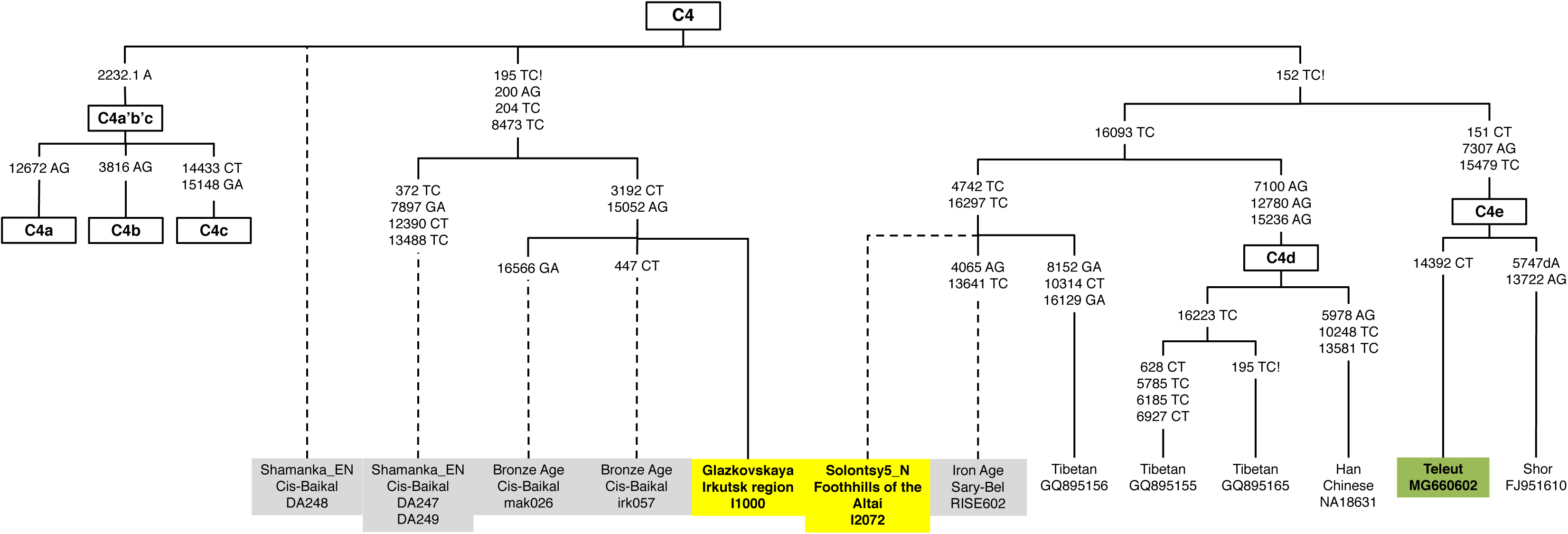
Schematic phylogeny of mtDNA haplocluster C4 sequences

#### Haplogroups C4a1 and C4a2

The Tofalar show high frequencies (>50%) of haplogroup C4a, a pattern observed in our previous study of the Tofalar, Udinsk Buryat, and Evenki [10]. In most cases, the results initially inferred from RFLP-analysis and HVSI sequencing have been confirmed and extended by subsequent studies (new and published) based on complete mitochondrial DNA analysis [11,12,28, the study herein]. Thus, C4a1 haplotypes, common in the Tofalar, Todzhi, Evenki, Even, and northern Altaian (Kumandin, Chelkan, and Teleut) samples, when assembled into one phylogenetic tree with their ancient counterparts, have redefined the ancestral C4a1 types. At its widest extent, the carriers of C4a1 lineages and sublineages spread from the Himalayas, Mongolia, and Iran across the south-central Siberia to Chukotka (Supplementary Fig. 2), contrasting with C4a2 diversity which is basically restricted to C4a2a (Supplementary Fig. 3).

#### Haplogroup C5

The phylogeny of C5 is structured into four major branches, assigned as C5a, C5b, C5c, and C5d, in accord with the latest release of PhyloTree (Supplementary Fig. 4-6). While the C5a1 and C5a2a samples are from the Altaic-speaking populations scattered across the southern extent of Eastern Siberia, the C5a2b is likely to reflect different migrations of Reindeer Koryak, Chukchi, Yukaghir, and Kamchatkan Even. An updated C5-m.16093T>C network, including 29 entire sequences, of which 15 are new, is given in Supplementary Fig. 4. Accordingly, the entire tree splits into two main lineages, C5c and C5d. While the C5c-m.16291C>T lineage is confined largely to the Teleut and Kumandin from southwestern Siberia, C5d1 is associated with the groups spread in the central/northern area where the Tungus-speaking populations expanded. In addition to Okunevo samples previously reported from West Siberia [42], ancient samples of this type were attested in the TransBaikal; one is Mesolithic (irk00x) and the other is pre-Bronze Age time depth (irk078) [43].

The tree of haplogroup C5b (Supplementary Fig. 6) is conspicuous for the C5b1a1 sublineage harbored by 4 of 39 (10.3%) Nganasan, 4 of 40 (10.0%) Tofalar, and 3 of 51 (5.9%) Todzhi individuals. The entire picture indicates a relatively recent affinity of the Nganasan from the Taimyr Peninsula to the Todzhi and Tofalar people. This finding is congruent with phylogenetic analysis of autosomal *HLA* class II genes in the respective populations. On all the neighbor-joining trees based on gene frequencies at *HLA* class II loci, the Nganasan cluster together with the Tofalar and Todzhi populations [43,45], thus corroborating a close genetic link between reindeer hunters of the Taimyr Peninsula and reindeer breeders of the eastern Sayan Mountains.

#### Haplogroup Z1

A novel Z1 sublineage, designated here as Z1b, and formed by two Tofalar samples, with one haplotype from western Sayan [10] and the other from eastern Sayan (this study) is noteworthy (Supplementary Fig. 7). An ancient sequence falling within the same sublineage of Z1, harboring new m.9804G>A, m.10295A>G, and m.15172G>A, was identified in the Korchugan site from the Novosibirk region, which must have arrived in Central Siberia by the Neolithic, recovering it from an individual dated to 5206-4805 calBCE (I0991). We also document an instance of Z1a1 in the nearly reported ancient sample I0998 (Khuzhir site from Olkhon Island in Lake Baikal, 2835-2472 calBCE). An entire tree features Z1 coalescing at ∼11.8 kya, whereas the Z1a node is much younger, ∼4.9 kya. Distinct sub-lineages of the Z1a haplogroup and their derivative haplotypes harbored by the Tubalar, Tofalar, Nganasan, Yukaghir, Koryak and Tungusic-speaking Evenki, Even, and Ulchi, could be a major part of Neolithic dispersals from southeastern Siberia, with Z1a1a emerged in the present-day Ket, Volga-Urals, and Finns and Saami much more recently. Occurrence of the Z1a1b haplotypes in single carriers, referred to previously as Nganasan or Yukaghir [11,12], in fact may be of Tungusic origin due to relatively recent gene flow from nearby Evenki or Even groups that patchily inhabited Arctic Siberia since early the period of Russian expansion.

#### Haplogroup F1b1

Of the 51 mt genomes sampled from Todzhi, who have long inhabited the refuge between western and estern Sayan Mountains, the territory of presumptive origin of some of the Ket’s ancestors, 8 individuals (15.7%) fall into two different sublineages of the F1b1b haplogroup (Supplementary Table 1, Supplementary Fig. 8). Accordingly, the phylogeny of F1b1b is structured into three major sub-branches, assigned as F1b1b1, F1b1b2, and F1b1b3. It is not surprising to observe the Ket identity of F1b1b2 confirmed by sharing of m.10227T>C and m.16179C>T variant, revealed earlier in 27.3% of Ket restricted to Podkamennaya Tungusska [21]. Remarkably, F1b1b2 was attested in a Late Bronze Age individual from Afontova Gora on the left bank of the Yenisei River (RISE553), and F1b1b3 in another, roughly contemporaneous sample from the same site and archaeological culture (RISE554). Recently, older samples (F1b1b) were attested with deeper, pre-Bronze Age time depth in the region west and north of Baikal Lake (irk036, irk068) [43].

## Discussion

### Postglacial recolonization of Northeastern Siberia

An unresolved issue in the postglacial resettlement of the Siberian Arctic is population substructure before the era of pastoralism, which emerged in the form of reindeer herding after ∼1200 CE [46] and could have had significant impacts on the population-genetic landscape of this vast region. The Taimyr Peninsula appear to be the last refuge for Nganasan, a palimpsest of reindeer hunters, the least touched genetically by surrounding herding groups [11, 47-49]. By the mid-17th century, the remnants of the westernmost Yukaghir tribe (Tavghi), the main ancestors of the Nganasan, was fused with the Yenisei Samoyeds (Somatu tribe) giving rise to the formation of the Avam Nganasan. A small subgroup of Yukagirized Tungus, later Vadey Nganasan, was recorded only in 1897 [6]. As recently as the 1950s, the Nganasan retained a lifestyle comparable to, and arguably continuous with that of pre-pastoral reindeer-hunters [46,50,51].

A major complication in studying genetic prehistory of Yukaghir tribes is profound admixture with NeoSiberians, the Tungus-speaking populations and (to lesser extent) the Yakut [11,52,53]. This admixture is a challenge for learning about the historical relationships amongst the Yukaghir and surrounding populations within a loose and poorly-defined North Siberian ethno-cultural complex composed of linguistically heterogeneous groups that, at the beginning of the Russian exploration, inhabited a region of northeastern Siberia, roughly between the Arctic Ocean in the north, the Verkhoyansk range in the west and southwest, the Kolyma range in the southeast, and the Pacific coast in the east [54, p.198]. To overcome this complication, we restricted analyses to mtDNAs attributed to Yukaghir-specific haplogroups, to C4b and D4b1c (D3), from adjoining groups, and to mtDNAs from more distant populations without evidence of admixture. The pattern of the basal branches of C4b detected among the modern and ancient inhabitants sampled from the Yana-Indigirka-Kolyma-Anadyr interfluvial (Supplementary Fig. 9), suggest that the initial episodes of the C4b diversification occurred in the Lena/Amga/Aldan area (south-central Yakutia) in the Neolithic and was then dispersed by geographic expansion during the Bronze Age and later. This conjecture is supported by an exceptionally large variety of C4b sublineages sharing C4b root (m.3816A>G) in populations of different linguistic affiliation. Interaction between reindeer Koryak, Chuvan, Khodyn and Anaul (the latter two tiny tribes dissolved among the Chuvan (Chuvantsi) shortly after the first Russians appeared in Chukotka in the mid-17^th^ century) is consistent with historical records indicating that quite a few of Yukaghir and Chuvantsi women were amongst the Koryak and the Chukchi by the end of 19^th^ century [6,55]. We also highlight the evolutionary history of D4b1c/D3, whose intrinsic diversity has not yet been well resolved. Following increased sampling, the haplogroup D4b1c (D3) phylogeny encompasses seven Nganasan and four Yukaghir, which together account for 61.1% of the entire D3 sample tested at the complete mtDNA level (Supplementary Table 1; Supplementary Fig. 10). Furthermore, D4b1c/D3 in samples irk022 (2455-2200 calBCE) of Bronze Age from Cis-Baikal, N5a (4340-4235 calBCE) of Middle Neolithic from Yakutia recently determined by Kılınç et al. (2018) [43], serve as additional support for the Nganasan/Yukaghir/Chuvan association. Co-representation of C4b and D3, being highly confident in the Neolithic mtDNA genomes sampled from Yakutia [43], suggest that the ancestry of the Yukaghir is multipartite and derived from the area that encompassed the East Sayan, Lena/Aldan valleys, and the northernwestern vicinity of Lake Baikal. The genetic continuity characterizing the ancestors of the Yukaghir was apparently interrupted by the arrival of a new population who arrived from Siberia taiga but whose original homeland was the in Trans-Baikal Mountains of north Manchuria [56,57,54, p.198].

## Conclusion

Here, we were assembled novel data on unique native Siberian populations, the Ket, Todzhi, and Tofalar. The timing and whereabouts of Yenisei Ket population history are long-standing issues [21,44,58]. The new mtDNA data provide a clearer picture of the number and timing of founding lineages. Our findings appear to reflect the origins and expansion history of mtDNA lineages that evolved in South-Central Siberia bordering Central-Inner Asia, as well as multiple phases of connections between this region and distant parts of Eurasia. Our results indicate that despite some level of continuity between ancient groups and present-day populations, the entire region exhibits a complex population history during the Holocene.

## Methods

### Mitochondrial genome sequencing for modern samples

Genomic DNA was extracted from blood buffy coats using standard procedures. The complete sequencing procedure of modern samples entailed PCR amplification of 2 overlapping mtDNA templates, which were sequenced with an Illumina HiSeq 2000 instrument [Illumina Human mtDNA Genome Guide 15037958B]. Short reads were analyzed with the BWA-backtrack tool [59] and Unipro UGENE version 1.21 software [60].

### Ancient DNA Analysis

In a dedicated clean room at Harvard Medical School, we prepared powder from the teeth of 7 individuals, all of whom we directly radiocarbon dated using accelerator mass spectrometry. These individuals consisted of:

- I0991 (Korchugan1-3, Novosibirsk, Korchugan 1; 5206-4805 calBCE (6060±50 BP, Poz-83427)),
- I0992 (Korchugan1-7, Novosibirsk, Korchugan 1; 5002-4730 calBCE (5990±50 BP, Poz-83428)),
- I0998 (Khuzhir-2, Irkutsk, Olkhon Island, Lake Baikal, Khuzhir; 2835-2472 calBCE (4040±35 BP, Poz-83426)),
- I1000 (Obkhoy-7, Irkutsk, Kachugskiy, Obkhoy; 2871-2497 calBCE (4100±40 BP, Poz-83436)),
- I2068 (230/3, Kurgan 2, Sayan Mountain, Minusinskaya Intermountain Basin, Tepsei III; 420-565 calCE (1560±30 BP, Poz-83507)),
- I2072 (230/13, Altai foothills, Solontcy-5; 3959-3715 calBCE (5050±40 BP, Poz-83497)),
- I2074 (230/18, Burial 1, Vas’kovo 4, Intermountain basin between spurs of Altai and Sayan Mountains; 5602-5376 calBCE (6520±40 BP, Poz-83514)). Generation of the ancient DNA followed previously established methodology,

so we do not discuss them in detail here [61]. Briefly, we extracted DNA [62], and converted it into individual barcoded double-stranded libraries in the presence of Uracil-DNA Glycosylase (UDG) and Endo VIII (USER, New England Biolabs) to characteristic errors at all but the final nucleotide [63]. We enriched the libraries for sequences overlapping the mitochondrial genome [64], and then sequenced on an Illumina NextSeq500 instrument using v.2 150 cycle kits for 2×76 cycles and 2×7 cycles. We computationally removed the barcode and adapter sequences, and merged pairs of reads requiring a 15 base pair overlap (allowing up to one mismatch). We mapped the merged sequeces to the reconstructed human mitochondrial DNA consensus sequence [65] using *bwa* (v.0.6.1) [58], removed sequences with the same strand orientation, start and stop positions, and built a consensus mitochondrial genome sequence for each sample (average coverages were 25-1550-fold). We used *contamMix-1.0.9* to estimate 95% confidence intervals for the rate of mismatch to the mitochondrial DNA consensus sequence, all of which were fully contained in the range 0.99-1 [66]. All samples had a rate of damage in the final nucleotide in the range of 0.028-0.127.

### Mitochondrial data analysis

All mtDNA genome consensus sequences were called using SAMTOOLS mpileup [67]. The resulting consensus sequences were then inspected by eye, with particular attention being paid to the hypervariable regions and nucleotide positions previously identified as being problematic [34]. All ambiguous sites were called as ‘N’. Mitochondrial haplogroups were assigned with mitohg v.0.2.8 software [68].

Entire mtDNA sequences were assembled into phylogenetic trees by using mtPhyl v5.003 software. Coalescence dates were estimated with the ρ statistic [69]. Standard errors (σ) were calculated according to Saillard et al. (2000) [70]. Mutational distances were converted into years using the substitution rate for the entire molecule, 2.67×10^−8^ substitutions per site per year [68]. The haplogroup affiliations reported in this analysis correspond to the current nomenclature of mtDNA in agreement with the latest release (February 2016) of PhyloTree Build 17 [34].

Overall, 218 newly reported mt genomes are listed in Table S1 along with ethnicity, sample location, and accession codes in GenBank. In the course of this study, 70 ancient mtDNA sequences were published and we used them to provide powerful new information about subhaplogroup affiliation (Supplementary Table 2) [24,25,30,33,41,43]. To uncover ancient mtDNA lineages, especially those related to the Altai-Sayan area, we compared modern mitogenomic data with their ancient counterparts sampled from skeletons recovered from the Euro-Siberian region that extends from the Central Europe and Scandinavia to Lake Baikal.

The sequence data for the new mitogenomes (n = 218) are available in Genbank with accession numbers MG660609-MG660750, MG778654-MG778683, MH807359, MH807364, MK180566, MK180567, MK180570, MK180572, MK180574, MK180577, MK180579, MK180580, MK217912-MK217947.

## General legend for phylogeny figures

In bold and green color is a new sequence generated through the course of this study. When two or more identical samples belong to the same group, their numbers are given in brackets. In bold and yellow are new ancient sequences generated through the course of this study, in grey - ancient sequences gleaned from published sources (see Supplementary Table 2). Dashed lines for ancient samples indicate that the sequence is not complete and contain gaps due to DNA damage or contamination.

## References

1. Pitulko, V., Pavlova, E. & Nikolskiy, P. Revising the archaeological record of the Upper Pleistocene Arctic Siberia: Human dispersal and adaptations in MIS 3 and 2. Quat. Sci. Rev. 165, 127–148 (2017).

2. Dolgikh, B. O. Rodovoi i plemennoi sostav narodov Sibiri v XVII veke. [The Clan and Tribal Composition of Siberian Peoples in the 17th century] (in Russian). (Akademia Nauk SSSR, 1960).

3. Levin, M. G. & Potapov, L. P. The Peoples of Siberia. (The University of Chicago Press, 1964).

4. Czaplicka, M. A. The Turks of Central Asia. (Clarendon Press, 1918).

5. Lopatin, I. A. The Extinct and Near-Extinct Tribes of Northeastern Asia as Compared with the American Indian. Am. Antiq. 5, 202–208 (1940).

6. Jochelson, W. I. The Yukaghir and the Yukaghirized Tungus. The Jesup North Pacific expedition, IX (1). (Memoirs of the AMNH) (E. J. Brill, 1910).

7. Torroni, A. et al. Asian affinities and continental radiation of the four founding Native American mtDNAs. Am. J. Hum. Genet. 53, 563–590 (1993a).

8. Torroni, A. et al. mtDNA variation of aboriginal Siberians reveals distinct genetic affinities with Native Americans. Am J Hum Genet 53, 591–608 (1993b).

9. Neel, J. V, Biggar, R. J. & Sukernik, R. I. Virologic and genetic studies relate Amerind origins to the indigenous people of the Mongolia/Manchuria/southeastern Siberia region. Proc. Natl. Acad. Sci. U. S. A. 91, 10737–41 (1994).

10. Starikovskaya, E. B. et al. Mitochondrial DNA diversity in indigenous populations of the southern extent of Siberia, and the origins of Native American haplogroups. Ann. Hum. Genet. 69, 67–89 (2005).

11. Volodko, N. V. et al. Mitochondrial Genome Diversity in Arctic Siberians, with Particular Reference to the Evolutionary History of Beringia and Pleistocenic Peopling of the Americas. Am. J. Hum. Genet. 82, 1084–1100 (2008).

12. Sukernik, R. I. et al. Mitochondrial genome diversity in the tubalar, even, and ulchi: Contribution to prehistory of native siberians and their affinities to native americans. Am. J. Phys. Anthropol. 148, 123–138 (2012).

13. Dryomov, S. V. et al. Mitochondrial genome diversity at the Bering Strait area highlights prehistoric human migrations from Siberia to northern North America. Eur. J. Hum. Genet. 23, 1399–1404 (2015).

14. Tackney, J. C. et al. Two contemporaneous mitogenomes from terminal Pleistocene burials in eastern Beringia. Proc. Natl. Acad. Sci. 112, 13833–13838 (2015).

15. Ruhlen, M. The origin of the Na-Dene. Proc. Natl. Acad. Sci. U. S. A. 95, 13994–13996 (1998).

16. Dolgikh, B. O. Kety [The Kets] (in Russian). (OGIZ, 1934).

17. Alekseenko, E. A. Kety: istoriko-demograficheskie ocherki [The Kets: Historical-ethnographic sketches] (in Russian). (Nauka, 1967).

18. Vainshtein, S. I. Tuvintsy-todzhintsy. Istorico-etnograficheskie ocherki [The Tuvan-Tozhu: Historical-ethnographic sketches] (in Russian). (Nauka, 1961).

19. Potapov, L. P. Etnicheskiy sostav i proiskhozhdeniye Altaitsev [Ethnic composition and origin of Altaians]. Istoriko-etnograficheskiy ocherk [Historical ethnographical essay] (in Russian). (Nauka, 1969).

20. Rassadin, V. I. The ethnic composition of Tofalar. 30, (2014).

21. Derbeneva, O. A., Starikovskaia, E. B., Volod’ko, N. V, Wallace, D. C. & Sukernik, R. I. Mitochondrial DNA variation in Kets and Nganasans and the early peoples of Northern Eurasia. Genet. 38, 1554–1560 (2002a).

22. Derbeneva, O. A., Starikovskaya, E. B., Wallace, D. C. & Sukernik, R. I. Traces of early Eurasians in the Mansi of northwest Siberia revealed by mitochondrial DNA analysis. Am. J. Hum. Genet. 70, 1009–1014 (2002b).

23. Sukernik, R. I., Volodko, N. V., Mazunin, I. O., Eltsov, N. P. & Starikovskaya, E. B. The genetic history of Russian old settlers of polar northeastern Siberia. Russ. J. Genet. 46, 1386–1394 (2010).

24. Allentoft, M. E. et al. Population genomics of Bronze Age Eurasia. Nature 522, 167–172 (2015).

25. Unterländer, M. et al. Ancestry and demography and descendants of Iron Age nomads of the Eurasian Steppe. Nat. Commun. 8, (2017).

26. Juras, A. et al. Diverse origin of mitochondrial lineages in Iron Age Black Sea Scythians. Sci. Rep. 7, 1–10 (2017).

27. Fernandes, V. et al. The Arabian cradle: Mitochondrial relicts of the first steps along the Southern route out of Africa. Am. J. Hum. Genet. 90, 347–355 (2012).

28. Derenko, M. et al. Complete mitochondrial DNA diversity in Iranians. PLoS One 8, (2013).

29. Naumova, O. I., Khaiat, S. S. & Rychkov, S. I. [Mitochondrial DNA diversity in Kazym Khanty] (in Russian). Genetika 45, 857–861 (2009).

30. Mathieson, I. et al. Genome-wide patterns of selection in 230 ancient Eurasians. Nature 528, 499–503 (2015).

31. Lamnidis, T. C. et al. Ancient Fennoscandian genomes reveal origin and spread of Siberian ancestry in Europe. Nat. Commun. 9, 5018 (2018).

32. Lazaridis, I. et al. Ancient human genomes suggest three ancestral populations for present-day Europeans. Nature 513, 409–413 (2014).

33. Günther, T. et al. Population genomics of Mesolithic Scandinavia: Investigating early postglacial migration routes and high-latitude adaptation. PLOS Biol. 16, e2003703 (2018).

34. van Oven, M. & Kayser, M. Updated comprehensive phylogenetic tree of global human mitochondrial DNA variation. Hum. Mutat. 30, 386–394 (2009).

35. Jeong, C. et al. The genetic history of admixture across inner Eurasia. Nat. Ecol. Evol. 3, 966–976 (2019).

36. Marrero, P., Abu-Amero, K. K., Larruga, J. M. & Cabrera, V. M. Carriers of human mitochondrial DNA macrohaplogroup M colonized India from southeastern Asia. BMC Evol. Biol. 16, 1–13 (2016).

37. Zhao, M. et al. Mitochondrial genome evidence reveals successful Late Paleolithic settlement on the Tibetan Plateau. Proc. Natl. Acad. Sci. 106, 21230–21235 (2009).

38. Qin, Z. et al. A mitochondrial revelation of early human migrations to the Tibetan Plateau before and after the last glacial maximum. Am. J. Phys. Anthropol. 143, 555–569 (2010).

39. Kashani, B. H. et al. Mitochondrial haplogroup C4c: A rare lineage entering America through the ice-free corridor? Am. J. Phys. Anthropol. 147, 35–39 (2012).

40. Potter, B. A. et al. Early colonization of Beringia and Northern North America: Chronology, routes, and adaptive strategies. Quat. Int. 444, 36–55 (2017).

41. Braje, T. J., Rick, T. C., Dillehay, T. D., Erlandson, J. M. & Klein, R. G. Arrival routes of first Americans uncertain-Response. Science 359, 1225 (2018).

42. Damgaard, P. de B. et al. The first horse hereders and the impact of early Bronze Age steppe expansions into Asia. Science. 360, eaar7711 (2018).

43. Kılınç, G. M. et al. Investigating Holocene human population history in North Asia using ancient mitogenomes. Sci. Rep. 8, 8969 (2018).

44. Uinuk-Ool, T. S., Takezaki, N., Derbeneva, O. A., Volodko, N. V & Sukernik, R. I. Variation of HLA class II genes in the Nganasan and Ket, two aboriginal Siberian populations. Eur. J. Immunogenet. Off. J. Br. Soc. Histocompat. Immunogenet. 31, 43–51 (2004).

45. Uinuk-Ool, T. S., Takezaki, N., Sukernik, R. I., Nagl, S. & Klein, J. Origin and affinities of indigenous Siberian populations as revealed by HLA class II gene frequencies. Hum. Genet. 110, 209–226 (2002).

46. Khlobystin, L. P. Drevnyaya istoriya Taimyrskogo Zapolyaria i voprosi formirovaniya kul’tur Severa Evrazii [Ancient History of Taimyr and the Formation of the North Eurasian Cultures]. (In Russian). (“Dmitry Bulanin”, 1998).

47. Dolgikh, B. O. Origin of the Nganasan. [Proishozhdenije nganasanov] (in Russian). Tr. instituta Etnogr. im. N. Mikluho-Maklaja AN SSSR. 3, 5–87 (1952).

48. Gol’tsova, T. V & Sukernik, R. I. [Genetic structure of an isolated group of the indigenous population of northern Siberia, the Nganasans (Tavginians) of Taimir] (in Russian). Genetika 15, 734–744 (1979).

49. Karaphet, T. M., Sukernik, R. I., Osipova, L. P. & Simchenko, Y. B. Blood groups, serum proteins, and red cell enzymes in the Nganasans (Tavghi)-reindeer hunters from Taimir Peninsula. Am. J. Phys. Anthropol. 56, 139–145 (1981).

50. Simchenko, Y. B. Kultura okhotnikov na oleney severnoy Evrazii. [The culture of reindeer hunters in northern Eurasia.] (in Russian). (Nauka, 1976).

51. Pitul’ko, V. Terminal Pleistocene—Early Holocene occupation in northeast Asia and the Zhokhov assemblage. Quat. Sci. Rev. 20, 267–275 (2001).

52. Fedorova, S. A. et al. Autosomal and uniparental portraits of the native populations of Sakha (Yakutia): Implications for the peopling of Northeast Eurasia. BMC Evol. Biol. 13, 127 (2013).

53. Duggan, A. T. et al. Investigating the prehistory of Tungusic peoples of Siberia and the Amur-Ussuri region with complete mtDNA genome sequences and Y-chromosomal markers. PLoS One 8, 1–19 (2013).

54. Zgusta, R. The Peoples of Northeast Asia through Time. Precolonial Ethnic and Cultural Processes along the Coast between Hokkaido and the Bering Strait. (Brill Academic Pub, 2015).

55. Jochelson, W. The Koryak. Material Culture and Social Organization of the Koryak. A publication of the Jesop North Pacific Expedition, X (2). (Memoir of the American Museum of Natural History). (E. J. Brill, 1908).

56. Okladnikov, A. P. Istoriya Jakutskoj ASSR [A history of the Yakut ASSR]. (in Russian). (AN SSSR, 1955).

57. Okladnikov, A. P. & Derevyanko, A. P. Dalekoye proshloye Primoriya i Priamuryia [The distant past of Primorie and Priamurie] (in Russian). (Dalnevostochnoye knizhnoye izdatel’stvo, 1973).

58. Flegontov, P. et al. Genomic study of the Ket: A Paleo-Eskimo-related ethnic group with significant ancient North Eurasian ancestry. Sci. Rep. 6, 1–12 (2016).

59. Li, H. & Durbin, R. Fast and accurate long-read alignment with Burrows-Wheeler transform. Bioinformatics 26, 589–595 (2010).

60. Okonechnikov, K. et al. Unipro UGENE: A unified bioinformatics toolkit. Bioinformatics 28, 1166–1167 (2012).

61. Haak, W. et al. Massive migration from the steppe was a source for Indo-European languages in Europe. Nature 522, 207–211 (2015).

62. Dabney, J. et al. Complete mitochondrial genome sequence of a Middle Pleistocene cave bear reconstructed from ultrashort DNA fragments. Proc. Natl. Acad. Sci. 110, 15758–15763 (2013).

63. Rohland, N., Harney, E., Mallick, S., Nordenfelt, S. & Reich, D. Partial uracil-DNA-glycosylase treatment for screening of ancient DNA. Philos. Trans. R. Soc. B Biol. Sci. 370, 20130624–20130624 (2014).

64. Maricic, T., Whitten, M. & Paabo, S. Multiplexed DNA sequence capture of mitochondrial genomes using PCR products. PLoS One 5, e14004 (2010).

65. Behar, D. M. et al. A ‘copernican’ reassessment of the human mitochondrial DNA tree from its root. Am. J. Hum. Genet. 90, 675–684 (2012).

66. Fu, Q. et al. A revised timescale for human evolution based on ancient mitochondrial genomes. Curr. Biol. 23, 553–559 (2013).

67. Li, H. et al. The Sequence Alignment/Map format and SAMtools. Bioinformatics 25, 2078–2079 (2009).

68. Dryomov, S. V. mitohg v.2.0.8. (2019). Available at: https://github.com/stasundr/gomitohg.

69. Forster, P., Harding, R., Torroni, A. & Bandelt, H.-J. Origin and evolution of Native American mtDNA variation: a reappraisal. Am. J. Hum. Genet. 59, 935–45 (1996).

70. Saillard, J., Forster, P., Lynnerup, N., Bandelt, H.-J. & Nørby, S. mtDNA Variation among Greenland Eskimos: The Edge of the Beringian Expansion. Am. J. Hum. Genet 67, 718–726 (2000).

