## Supplemental materials for "Mitochondrial Genome Diversity in the Central Siberian Plateau with Particular Reference to Prehistory of Northernmost Eurasia"

#### Samples from South Siberian archaeological sites

##### Korchugan-1

Human fossils from the Neolithic cemetery located in the pre-taiga zone of the West Siberian Plain (Kyshtovka Region of Novosibirsk Province; N 56°28′; E 76°18′). Koshkurgan-1, a burial yard with earthen mounds, were excavated by Academician V.I. Molodin in 1996. Available calibrated radiocarbon dates indicate an age of 6$^{th}$ – 5$^{th}$ centuries BCE. Three tombs were attributed to the Neolithic period; human fossils from burials 3 and 7 are the focus of the work here. The three Neolithic tombs form a row oriented along the NW-SE line. The complete information is provided in the paper "The Neolithic burial ground of Korchugan in the Middle Tara" by V.I. Molodin, A.V. Novikov, T.A. Chikisheva in the book "Neolithic-Charcolithic Period in southern West Siberia. Kemerovo: Izdat. Kuzbassvuzizdat, 1999, pp. 66 – 98."

Burial 3 was established in an ovoid grave, 2.2 x 0.75 x 0.25-0.17 m in size; the grave floor was uneven with one of the short sides higher than the opposite one. The occupant is an adult male of 45 – 50 years of age, large in stature, and the features of the bones testify to well-developed muscles. The body was placed in the tomb in the stretched supine position with the head towards N-NE.

Burial 7 was established in the grave, 2.23 x 1.05 x 0.5-0.64 m in size. The occupant was estimated as a female 17-23 years of age. The body was placed in the tomb in the stretched supine position with the head toward the N. The hands were placed over the hip bones. Pendants of squirrel (?) canine teeth were found close to the right knee. A necklace of nine miniature flat bird figurines made of bone with hanging openings was found over the chest. The tenth element in the necklace was a tear-shaped bone pendant.

Craniometric and the long bone measurement data from the Neolithic burials at Korchugan-1 are provided in the abovementioned paper.

The features of Korchugan-1 burial rite are similar to those noted at the Neolithic sites in the Ob basin and likely represent a single historical-cultural unity. However, the area was populated by anthropologically diverse populations. The Korchugan-1 materials represent an anthropologically distinct group related to the population of the Altai Plain and Kuznetsk Hollow.

##### I2072 (230/13 – Solontsy-5, tomb1)

Tomb 1 was found in the earthen mound yard of Solontsy-5 on the terrace remnant on left side of the Biya at the confluence with the Chepshushka: N 52°29'; E 86°13'.

The site is situated in the forest-steppe zone of the Altai-Salair piedmonts. Excavations were carried out in 2000 – 2001 by Natalia Y. Kungurova. The age of the site was estimated as 4850–4502 BCE, and the burial features attest the Kuznetsk-Altai archaeological culture. All the nine tombs of the site were oriented along the NW – SE line. The dead were placed in the tombs in the stretched supine position, the arms were stretched along the body, the heads were oriented towards the NE-E.

The sample represents the upper medial canine tooth of an adult individual – male (?) of an estimated age of 35 – 40 years. Remains of an infant, which age was estimated to approximately 18 months basing on the dental system features and the size of the femurs was found in the same tomb. The skull is poorly preserved due to the post-mortem deformation. The skull is moderately developed, and the mandible is relatively gracile. The estimated stature varies from 161.7 cm to 166.5 cm depending on the technique.

**I2074 (230/18 – Vaskovo-4, tomb 1)**

The Vaskovo-4 (N 55°03′; E 85°05′) is an earthen burial mound yard located on the high right bank of the Tykhta close to its junction with the Inia in the vicinity to the village of Vaskovo. The cemetery was studied by Y.M. Borodkin in 1967; tomb 1 was excavated by V.V. Bobrov in 1979. The burial features attest the Kuznetsk-Altai archaeological culture. The deceased was placed in the tomb in the supine position with the head towards N – NE. The site is situated in the Kuznetsk Hollow in the forest-steppe zone of the Altai-Salair piedmonts.

Osteological analysis indicates an adult male (Maturus). The sample for genetic analysis represents the second upper premolar.

**I2068 (230/3 – Tepsei-3, mound 2)**

A cluster of cemeteries of various historical-cultural periods (N 53.96; E 91.56) was situated at the Tepsei Mount on the Yenisei right-side bank at the confluence with the Tuba at 25 km northwards from Minusinsk. Rescue archaeological work was carried out at the site under the project of the Sayan-Shushenski Electric Power Station construction in 1968 – 1970. The burial sites were attributed to the Afanasievo culture of the Charcolithic Period until the Yenisei Kyrgyz Culture of the early medieval period. Mound 2 was initially attributed to the Afanasievo Culture; the features of the burial rite were not published. The radiocarbon date of 420-565 calCE (1560±30 BP, Poz-83507) indicate the period of the Tashtyk Culture.

The sample for genetic analysis represents the upper molar of an adult male of 30 – 40 years old.

**I0998 (Khuzhir-2, tomb 2)**

The tomb was located in the northwestern part of the Olkhon Island in Lake Baikal (N 53.19; E 107.34). The site was excavated by A.P. Okladnikov in 1992. The grave (280 cm long and 65 cm wide) was covered with stone pavement, overlain by soil and turf. The tomb was damaged by looters. The bones of a male of 35 – 40 years old were scattered over the grave. The hands and feet were in situ indicating the stretched supine position of the dead. The site was attributed to the Serovo Culture. The sample for genetic analysis represents the second upper premolar.

**I1000 (Obkhoi, tomb 7)**

The burial yard was located close to the village of Obkhoi on the northwestern bank of Lake Baikal (N 54.02; E 105.47). Tomb 7 was excavated by A.P. Okladnikov in 1971. Human remains indicating a young male of 30 – 35 years old were found in the stretched supine position with the head towards the NW. The burial was attributed to the Glazkovo Culture. The sample for genetic analysis represents the second upper premolar.

**General legend for phylogeny figures:** In bold and green color is a new sequence generated through the course of this study. When two or more identical samples belong to the same group, their numbers are given in brackets. In bold and yellow are new ancient sequences generated through the course of this study, in grey - ancient sequences gleaned from published sources (see Supplementary Table 2). Dashed lines for ancient samples indicate that the sequence is not complete and contain gaps due to DNA damage or contamination.

Supplementary Figure 1. Phylogeny of M8a sequences.

Supplementary Figure 2. Phylogeny of C4a1 sequences.

Supplementary Figure 3. Phylogeny of C4a2a sequences.

Supplementary Figure 4. The phylogeny of C5 (C5c and C5d) sequences.

Supplementary Figure 5. The phylogeny of C5a sequences.

Supplementary Figure 6. The phylogeny of C5b sequences.

Supplementary Figure 7. The phylogeny of Z1 sequences.

Supplementary Figure 8. The phylogeny of F1b1 sequences.

Supplementary Figure 9. Updated phylogeny of C4b and its derivatives.

Supplementary Figure 10. Updated phylogeny D4b1c-D3 and its derivatives.

**Supplementary Figure 1.**

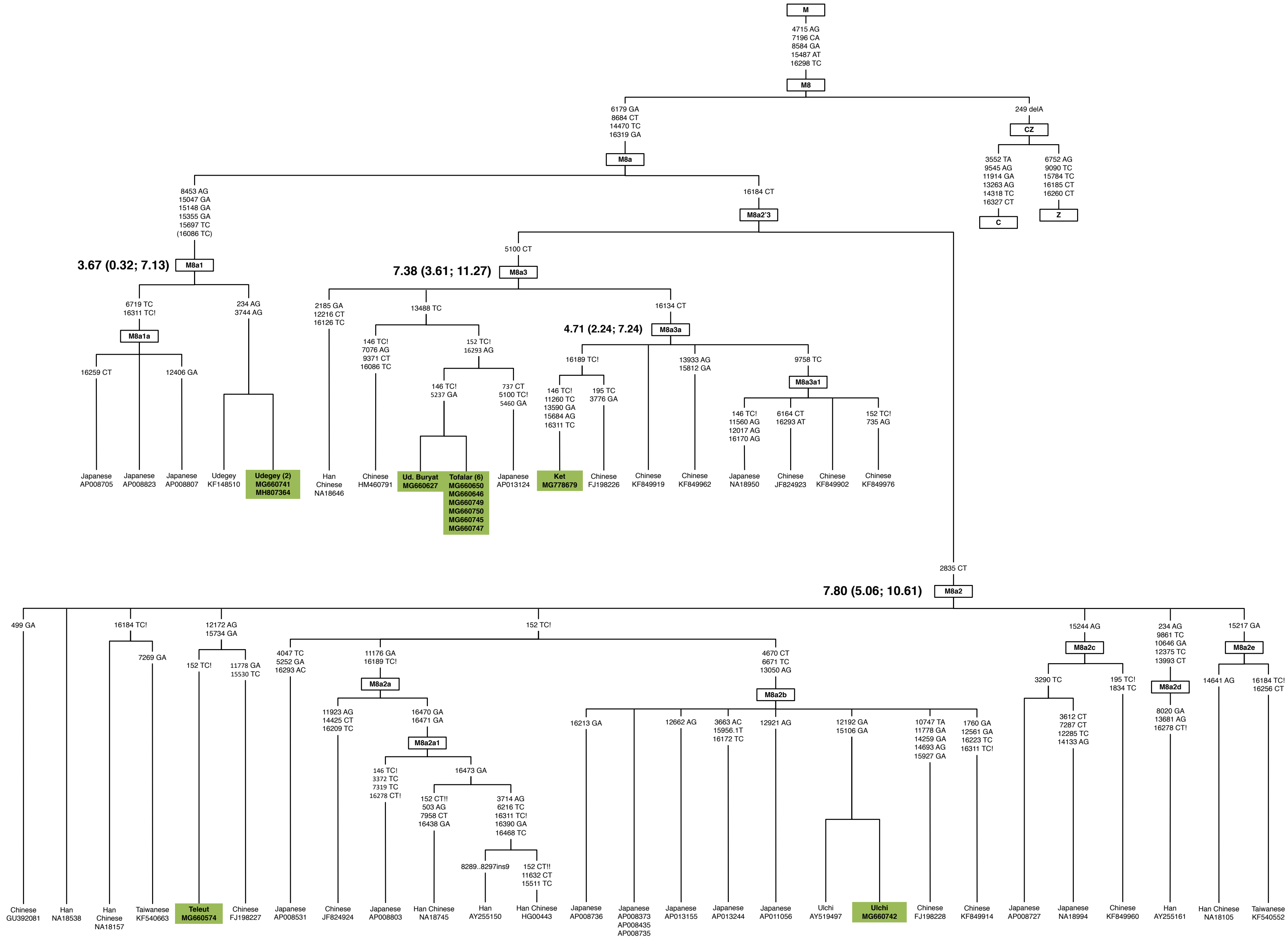

### Supplementary Figure 2.

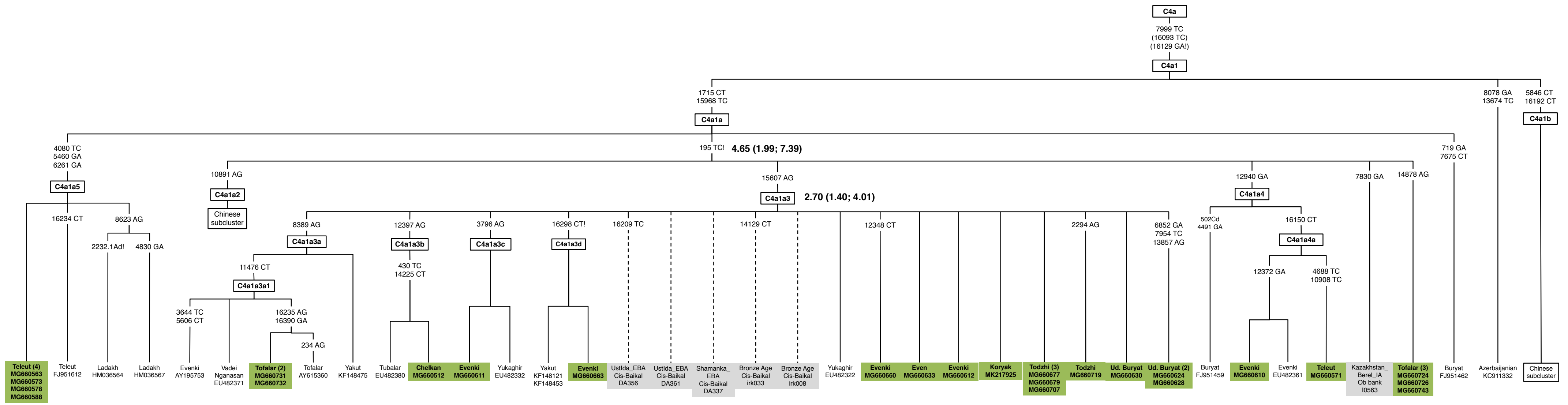

**Supplementary Figure 3.**

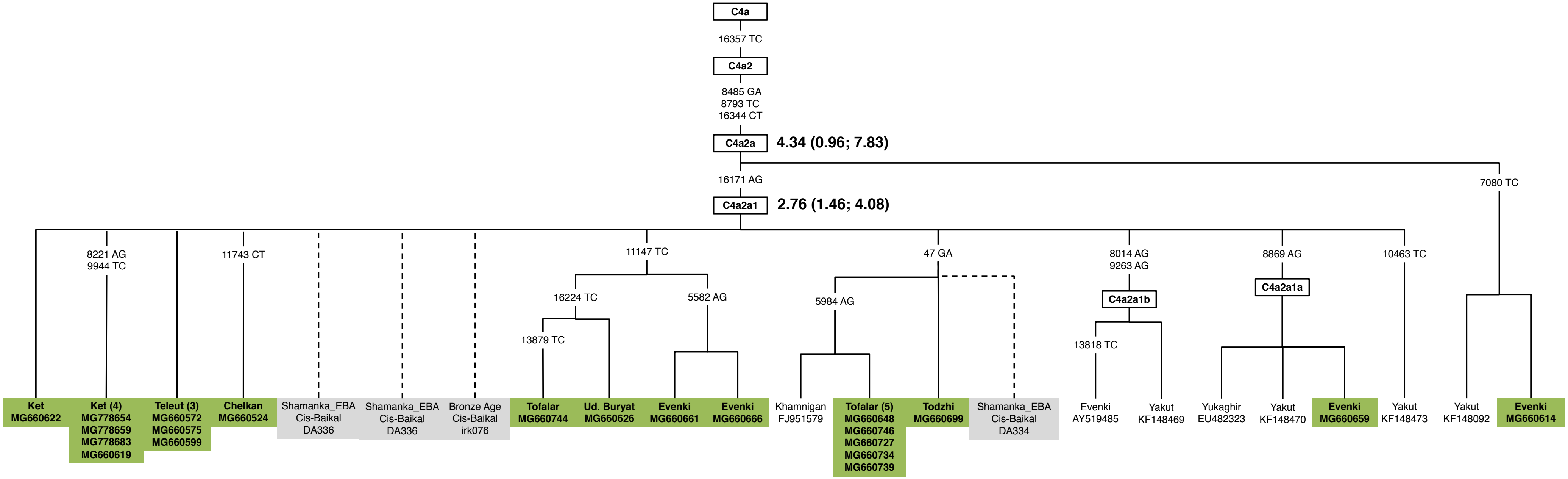

### Supplementary Figure 4.

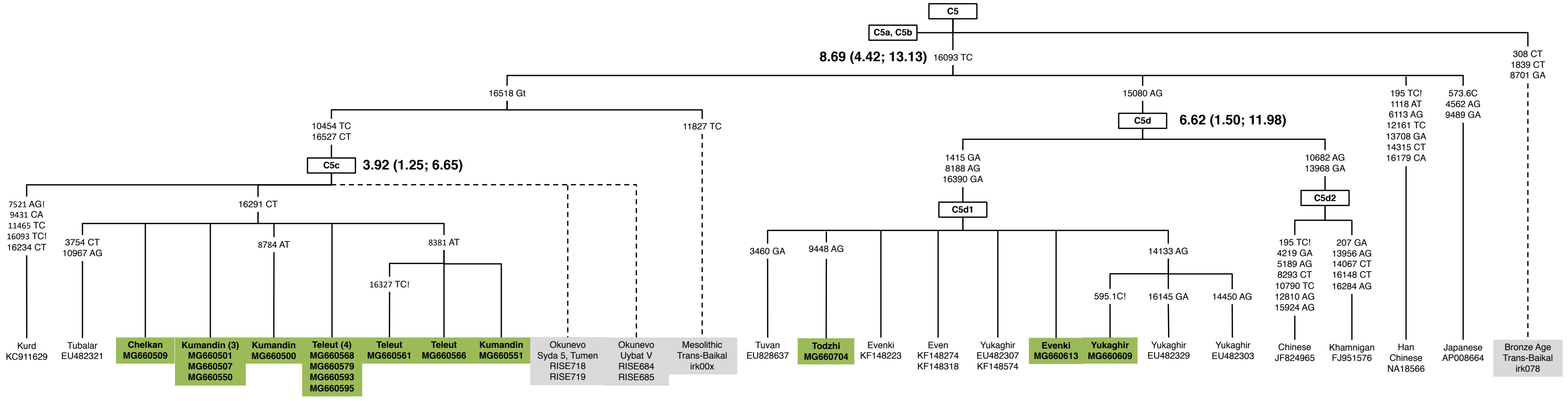

**Supplementary Figure 5.**

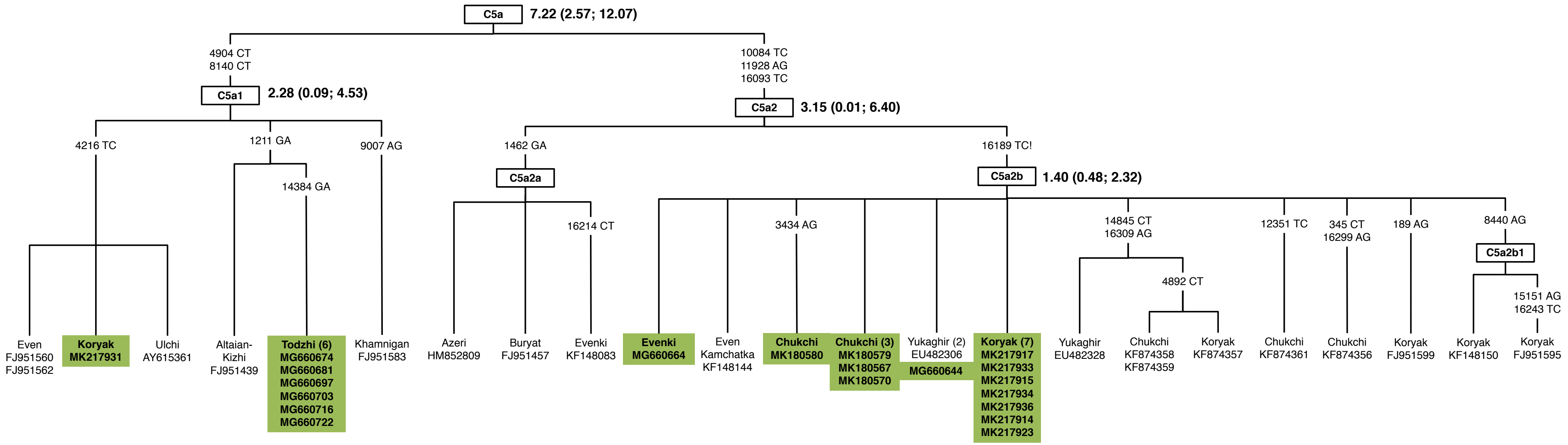

**Supplementary Figure 6.**

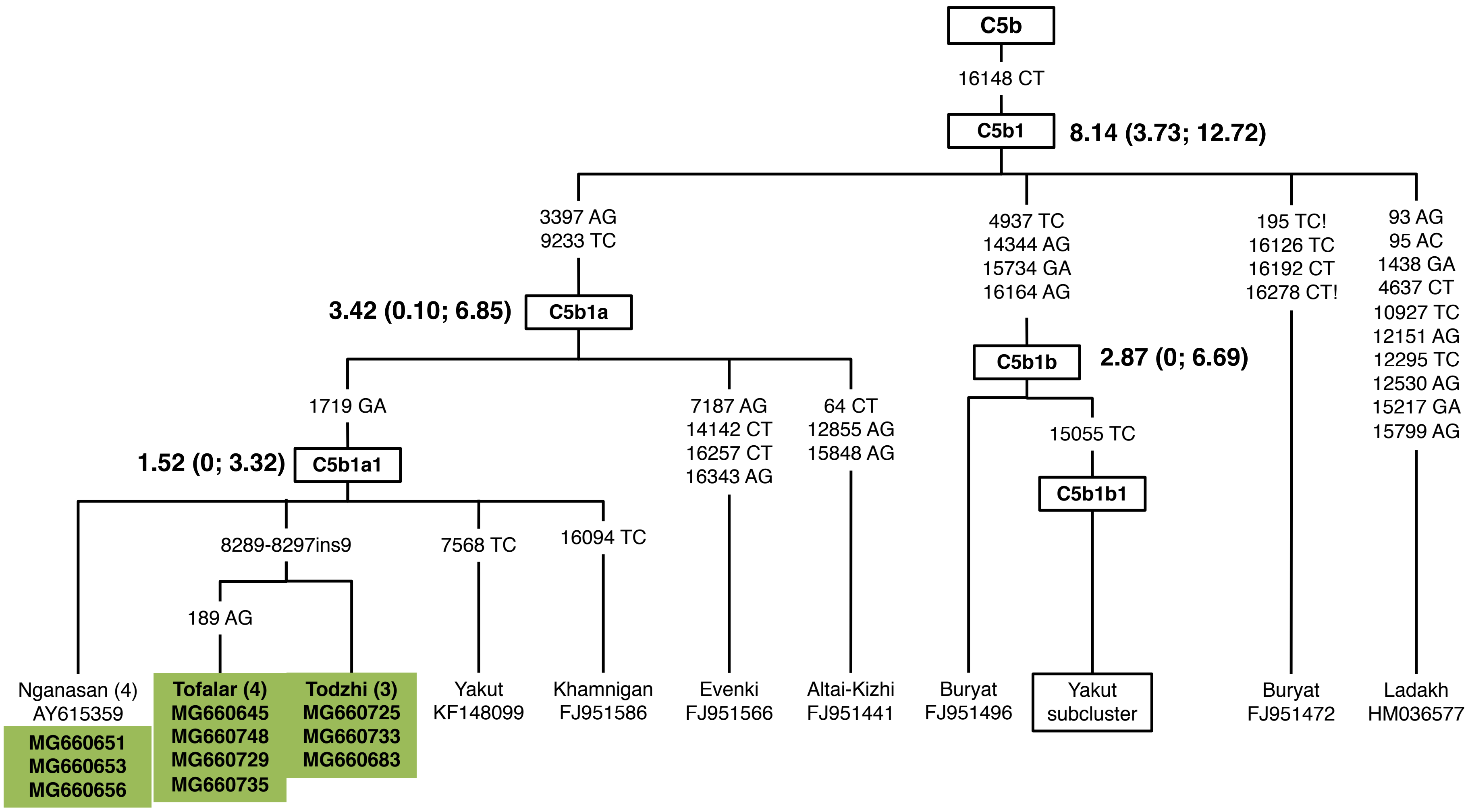

#### Supplementary Figure 7.

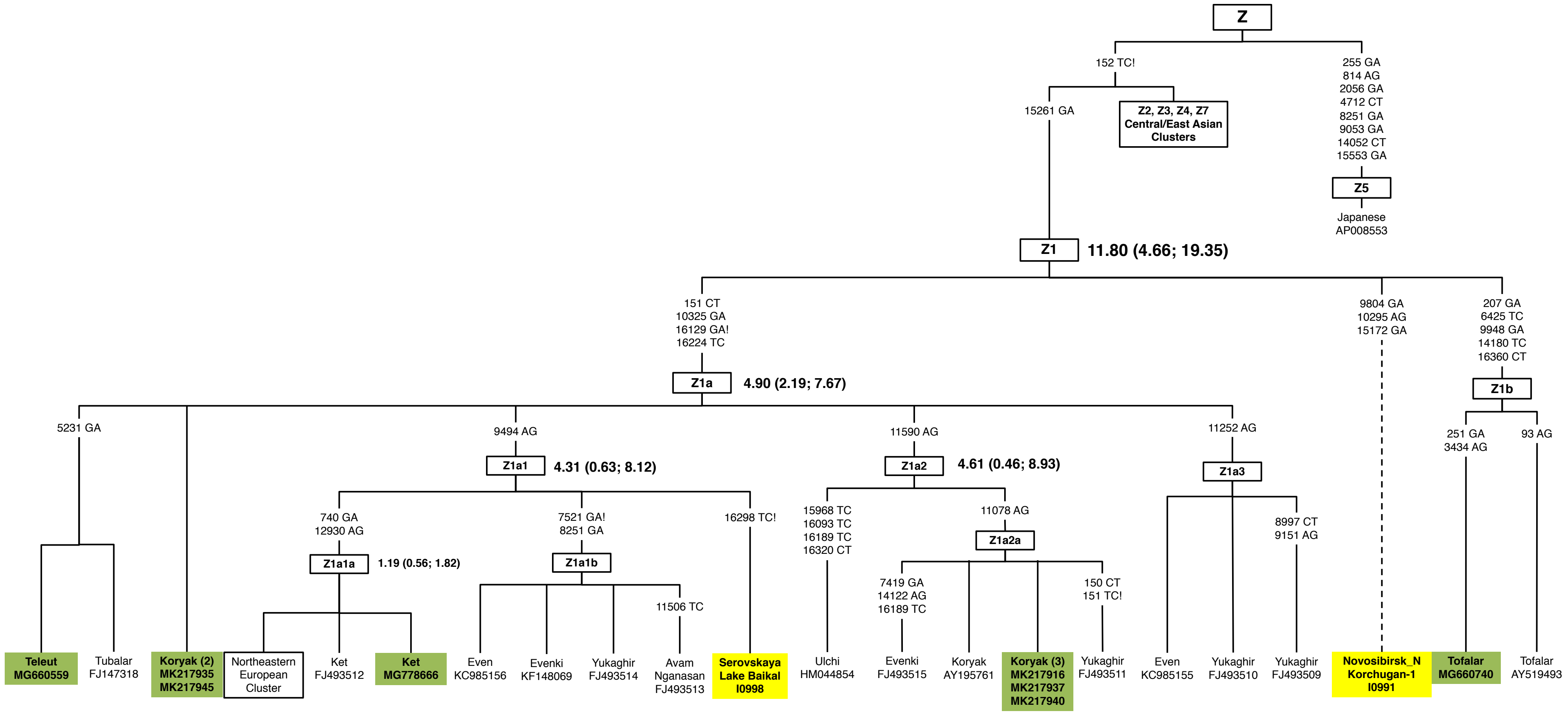

### Supplementary Figure 8.

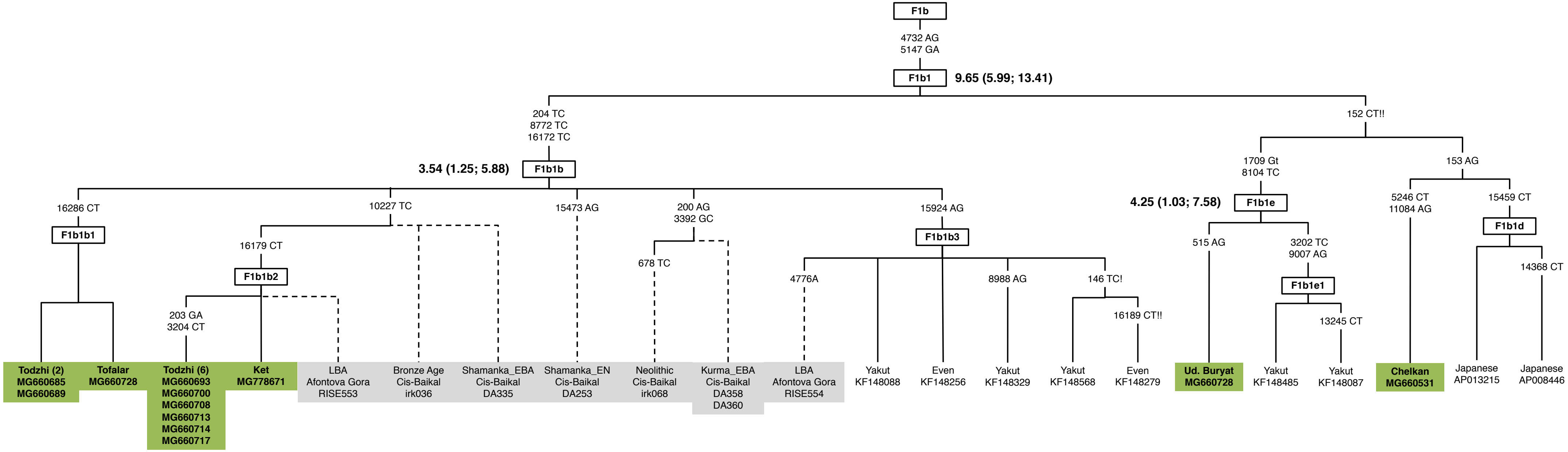

### Supplementary Figure 9.

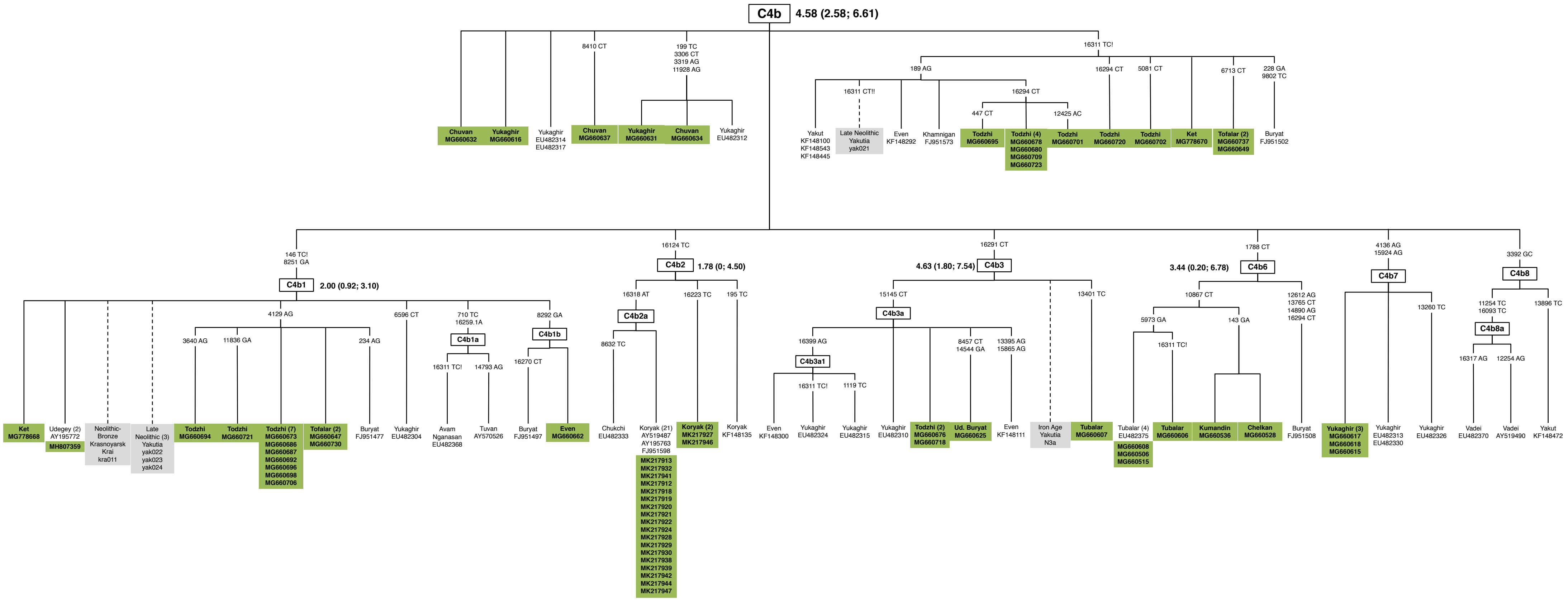

**Supplementary Figure 10.**

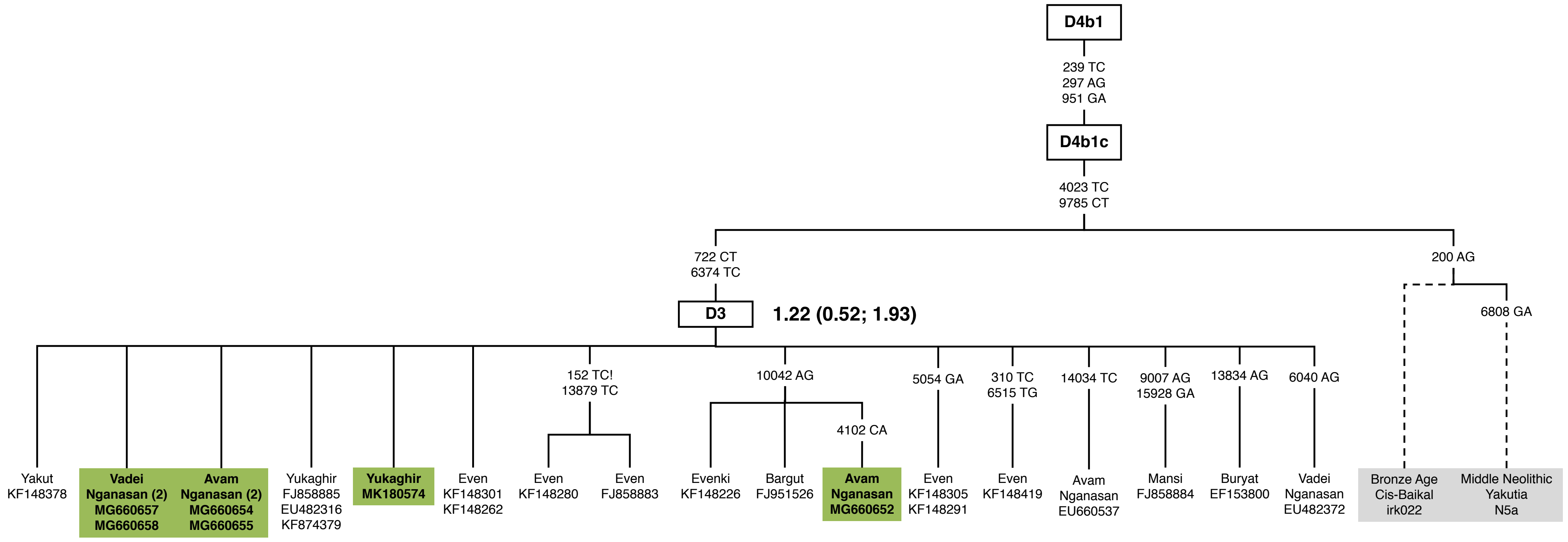

**Supplementary Table 1.** List of modern mtDNA samples generated in course of this study.

| # | Haplogroup | Ethnicity | Sampling location | Genbank ID | Distinguishing / Private mutations |
|---|---|---|---|---|---|
| 1 | A10 | Mansi | Shaim | MG660641 | C64T, G2831A, A16230G!, C16256T |
| 2 | A14 | Tuvan | Bai-Tal | MG660670 | A4123G, A8396G, T11253C, T16086C |
| 3 | A8a2 | Ket | Kellog | MG778662 | G235A!, A9007G, T15229C |
| 4 | A8a2 | Tuvan | Bai-Tal | MG660672 | A3912G, G7762A, T16223C |
| 5 | A8a2 | Tofalar | Alygdzher | MG660738 | G235A!, T15229C |
| 6 | A8b | Koryak | Karaginskiy district | MK217926 | - |
| 7 | B4a1c2 | Todzhi | Iiy | MG660688 | A16183C |
| 8 | B4j | Todzhi | Toora-Hem | MG660675 | 8281del10!, A15172G!, A16183C |
| 9 | C4a1a-195! | Tofalar | Alygdzher | MG660724 | A14878G |
| 10 | C4a1a-195! | Tofalar | Alygdzher | MG660726 | A14878G |
| 11 | C4a1a-195! | Tofalar | V. Gutara | MG660743 | A14878G |
| 12 | C4a1a3 | Evenki | Chumikan | MG660612 | - |
| 13 | C4a1a3 | Koryak | Karaginskiy district | MK217925 | - |
| 14 | C4a1a3 | Ud. Buryat | Kushun | MG660624 | G6852A, T7954C, A13857G, (T16093C) |
| 15 | C4a1a3 | Ud. Buryat | Kushun | MG660628 | G6852A, T7954C, A13857G |
| 16 | C4a1a3 | Ud. Buryat | Kushun | MG660630 | - |
| 17 | C4a1a3 | Even | Markovo | MG660633 | - |
| 18 | C4a1a3 | Evenki | Nelkan, Dzhigda | MG660660 | C12348T |
| 19 | C4a1a3 | Todzhi | Toora-Hem | MG660677 | - |
| 20 | C4a1a3 | Todzhi | Toora-Hem | MG660679 | - |
| 21 | C4a1a3 | Todzhi | Adir-Kezhig | MG660707 | - |
| 22 | C4a1a3 | Todzhi | Adir-Kezhig | MG660719 | A2294G |
| 23 | C4a1a3a1 | Tofalar | Alygdzher | MG660731 | A16235G, G16390A |
| 24 | C4a1a3a1 | Tofalar | Alygdzher | MG660732 | A16235G, G16390A |
| 25 | C4a1a3c | Evenki | Chumikan | MG660611 | - |
| 26 | C4a1a3d | Evenki | Nelkan, Dzhigda | MG660663 | - |
| 27 | C4a1a4a | Evenki | Chumikan | MG660610 | G12372A |
| 28 | C4a2a | Evenki | Chumikan | MG660614 | T7080C |
| 29 | C4a2a1 | Ket | Kellog | MG778654 | A8221G, T9944C |
| 30 | C4a2a1 | Evenki | Kellog | MG778683 | A8221G, T9944C |
| 31 | C4a2a1 | Ket | Kellog | MG778659 | A8221G, T9944C |
| 32 | C4a2a1 | Ket | Selivanikha | MG660619 | A8221G, T9944C |
| 33 | C4a2a1 | Ket | Bakhta | MG660622 | - |
| 34 | C4a2a1 | Ud. Buryat | Kushun | MG660626 | T11147C, T16224C |
| 35 | C4a2a1 | Tofalar | Nerkha | MG660648 | G47A, A5984G |
| 36 | C4a2a1 | Evenki | Nelkan, Dzhigda | MG660661 | A5582G, T11147C |
| 37 | C4a2a1 | Evenki | Val | MG660666 | A5582G, T11147C |
| 38 | C4a2a1 | Todzhi | Iiy | MG660699 | G47A |
| 39 | C4a2a1 | Tofalar | Alygdzher | MG660727 | G47A, A5984G |
| 40 | C4a2a1 | Tofalar | Alygdzher | MG660734 | G47A, A5984G |
| 41 | C4a2a1 | Tofalar | Alygdzher | MG660739 | G47A, A5984G |
| 42 | C4a2a1 | Tofalar | V. Gutara | MG660744 | T11147C, T13879C, T16224C |
| 43 | C4a2a1 | Tofalar | V. Gutara | MG660746 | G47A, A5984G |
| 44 | C4a2a1a | Evenki | Nelkan, Dzhigda | MG660659 | - |
| 45 | C4b | Yukaghir | Andrushkino | MG660616 | - |
| 46 | C4b | Yukaghir/ Chuvan | Markovo | MG660631 | T199C, C3306T, A3319G, A11928G |
| 47 | C4b | Yukaghir/ Chuvan | Markovo | MG660632 | - |
| 48 | C4b | Yukaghir/ Chuvan | Markovo | MG660634 | T199C, C3306T, A3319G, A11928G |
| 49 | C4b | Yukaghir/ Chuvan | Markovo | MG660635 | C8410T |
| 50 | C4b | Yukaghir/ Chuvan | Markovo | MG660636 | - |
| 51 | C4b | Yukaghir/ Chuvan | Markovo | MG660637 | C8410T |
| 52 | C4b-16311! | Evenki | Kellog | MG778670 | T16311C! |
| 53 | C4b-16311! | Tofalar | Nerkha | MG660649 | C6713T, T16311C! |
| 54 | C4b-16311! | Todzhi | Toora-Hem | MG660678 | A189G, C16294T, T16311C! |
| 55 | C4b-16311! | Todzhi | Toora-Hem | MG660680 | A189G, C16294T, T16311C! |
| 56 | C4b-16311! | Todzhi | Iiy | MG660695 | A189G, C447T, C16294T, T16311C! |
| 57 | C4b-16311! | Todzhi | Iiy | MG660701 | A189G, A12425C, C16294T, T16311C! |
| 58 | C4b-16311! | Todzhi | Iiy | MG660702 | T5081C, T16311C! |
| 59 | C4b-16311! | Todzhi | Adir-Kezhig | MG660709 | A189G, C16294T, T16311C! |
| 60 | C4b-16311! | Todzhi | Adir-Kezhig | MG660720 | C16294T, T16311C! |
| 61 | C4b-16311! | Todzhi | Adir-Kezhig | MG660723 | A189G, C16294T, T16311C! |
| 62 | C4b-16311! | Tofalar | Alygdzher | MG660737 | C6713T, T16311C! |
| 63 | C4b1 | Evenki/Selkup | Krasnoselkup | MG778668 | - |
| 64 | C4b1 | Tofalar | Nerkha | MG660647 | A4129G |
| 65 | C4b1 | Todzhi | Toora-Hem | MG660673 | A4129G |
| 66 | C4b1 | Todzhi | Iiy | MG660686 | A4129G |
| 67 | C4b1 | Todzhi | Iiy | MG660687 | A4129G |
| 68 | C4b1 | Todzhi | Iiy | MG660692 | A4129G |
| 69 | C4b1 | Todzhi | Iiy | MG660694 | G3640A, A4129G |
| 70 | C4b1 | Todzhi | Iiy | MG660696 | A4129G |
| 71 | C4b1 | Todzhi | Iiy | MG660698 | A4129G |
| 72 | C4b1 | Todzhi | Iiy | MG660706 | A4129G |
| 73 | C4b1 | Todzhi | Adir-Kezhig | MG660721 | A4129G, A11836G |
| 74 | C4b1 | Tofalar | Alygdzher | MG660730 | A4129G |
| 75 | C4b1 | Udegey | Agzu/Ternei | MH807359 | - |
| 76 | C4b1b | Evenki | Nelkan, Dzhigda | MG660662 | - |
| 77 | C4b2 | Koryak | Karaginskiy district | MK217927 | T16223C |
| 78 | C4b2 | Koryak | Karaginskiy district | MK217946 | T16223C |
| 79 | C4b2a | Koryak | Karaginskiy district | MK217912 | - |
| 80 | C4b2a | Koryak | Karaginskiy district | MK217913 | - |
| 81 | C4b2a | Koryak | Karaginskiy district | MK217918 | - |
| 82 | C4b2a | Koryak | Karaginskiy district | MK217919 | - |
| 83 | C4b2a | Koryak | Karaginskiy district | MK217920 | - |
| 84 | C4b2a | Koryak | Karaginskiy district | MK217921 | - |
| 85 | C4b2a | Koryak | Karaginskiy district | MK217922 | - |
| 86 | C4b2a | Koryak | Karaginskiy district | MK217924 | - |
| 87 | C4b2a | Koryak | Karaginskiy district | MK217928 | - |
| 88 | C4b2a | Koryak | Karaginskiy district | MK217929 | - |
| 89 | C4b2a | Koryak | Karaginskiy district | MK217930 | - |

| | | | | | |
|---|---|---|---|---|---|
| 90 | C4b2a | Koryak | Karaginskiy district | MK217932 | - |
| 91 | C4b2a | Koryak | Karaginskiy district | MK217938 | - |
| 92 | C4b2a | Koryak | Karaginskiy district | MK217939 | - |
| 93 | C4b2a | Koryak | Karaginskiy district | MK217941 | - |
| 94 | C4b2a | Koryak | Karaginskiy district | MK217942 | - |
| 95 | C4b2a | Koryak | Karaginskiy district | MK217944 | - |
| 96 | C4b2a | Koryak | Karaginskiy district | MK217947 | - |
| 97 | C4b3a | Ud. Buryat | Kushun | MG660625 | C8457T, C9028A(hp), G14544A |
| 98 | C4b3a | Todzhi | Toora-Hem | MG660676 | - |
| 99 | C4b3a | Todzhi | Adir-Kezhig | MG660718 | - |
| 100 | C4b7 | Yukaghir | Andrushkino | MG660615 | - |
| 101 | C4b7 | Yukaghir | Andrushkino | MG660617 | - |
| 102 | C4b7 | Yukaghir | Andrushkino | MG660618 | - |
| 103 | C5a1 | Koryak | Karaginskiy district | MK217931 | T4216C |
| 104 | C5a1 | Todzhi | Toora-Hem | MG660674 | G1211A, G14384A |
| 105 | C5a1 | Todzhi | Toora-Hem | MG660681 | G1211A, G14384A |
| 106 | C5a1 | Todzhi | Iiy | MG660697 | G1211A, G14384A |
| 107 | C5a1 | Todzhi | Iiy | MG660703 | G1211A, G14384A |
| 108 | C5a1 | Todzhi | Adir-Kezhig | MG660716 | G1211A, G14384A |
| 109 | C5a1 | Todzhi | Adir-Kezhig | MG660722 | G1211A, G14384A |
| 110 | C5a2b | Chukchi | Ayon | MK180567 | - |
| 111 | C5a2b | Chukchi | Ayon | MK180570 | - |
| 112 | C5a2b | Chukchi | Ayon | MK180579 | - |
| 113 | C5a2b | Chukchi | Ayon | MK180580 | A3434G |
| 114 | C5a2b | Koryak | Karaginskiy district | MK217915 | - |
| 115 | C5a2b | Koryak | Karaginskiy district | MK217917 | - |
| 116 | C5a2b | Koryak | Karaginskiy district | MK217933 | - |
| 117 | C5a2b | Koryak | Karaginskiy district | MK217934 | - |
| 118 | C5a2b | Koryak | Karaginskiy district | MK217936 | - |
| 119 | C5a2b | Koryak | Karaginskiy district | MK217914 | - |
| 120 | C5a2b | Koryak | Karaginskiy district | MK217923 | - |
| 121 | C5a2b | Yukaghir | Nelemnoye | MG660644 | - |
| 122 | C5a2b | Evenki | Nelkan, Dzhigda | MG660664 | - |
| 123 | C5b1a | Tofalar | Nerkha | MG660645 | A189G, 8289-8297ins9 |
| 124 | C5b1a | Avam Nganasan | Ust'-Avam | MG660651 | - |
| 125 | C5b1a | Avam Nganasan | Ust'-Avam | MG660653 | - |
| 126 | C5b1a | Avam Nganasan | Ust'-Avam | MG660656 | - |
| 127 | C5b1a | Todzhi | Toora-Hem | MG660683 | 8289-8297ins9 |
| 128 | C5b1a | Todzhi-Tuva | Alygdzher | MG660725 | 8289-8297ins9 |
| 129 | C5b1a | Tofalar | Alygdzher | MG660729 | A189G, 8289-8297ins9 |
| 130 | C5b1a | Todzhi-Tuva | Alygdzher | MG660733 | 8289-8297ins9 |
| 131 | C5b1a | Tofalar | Alygdzher | MG660735 | A189G, 8289-8297ins9 |
| 132 | C5b1a | Tofalar | V. Gutara | MG660748 | A189G, 8289-8297ins9 |
| 133 | C5d1 | Yukaghir | Cherskiy | MG660609 | A14133G |
| 134 | C5d1 | Evenki | Chumikan | MG660613 | - |
| 135 | C5d1 | Todzhi | Iiy | MG660704 | A9448G |
| 136 | D3 | Yukaghir | Yanranay | MK180574 | - |
| 137 | D3 | Avam Nganasan | Ust'-Avam | MG660652 | C4102A, A10042G |
| 138 | D3 | Avam Nganasan | Ust'-Avam | MG660654 | - |
| 139 | D3 | Avam Nganasan | Ust'-Avam | MG660655 | - |
| 140 | D3 | Vadei Nganasan | Factoriya Novaya, Khatanga | MG660657 | - |
| 141 | D3 | Vadei Nganasan | Factoriya Novaya, Khatanga | MG660658 | - |
| 142 | D4b1a2a1 | Koryak | Karaginskiy district | MK217943 | T11383C, A14122C, T16093C, C16519T |
| 143 | D4b1a2a1 | Chukchi | Ayon | MK180572 | T11383C, G11914A!, A14122C, T16093C, C16519T |
| 144 | D4b1a2a1 | Chukchi | Ayon | MK180566 | T11383C, A14122C, C15370T, T16093C, C16519T |
| 145 | D4b1a2a1 | Chukchi | Yanranay | MK180577 | T11383C, G11914A!, A14122C, T16093C, C16519T |
| 146 | D4b1a2a2 | Tuvan | Bai-Tal | MG660671 | G14207A, C16519T |
| 147 | D4b1a2a2 | Todzhi | Iiy | MG660690 | - |
| 148 | D4b1a2a2 | Todzhi | Iiy | MG660691 | - |
| 149 | D4j4 | Evenki | Nelkan, Dzhigda | MG660665 | T14470C |
| 150 | D4j4a | Evenki/Ulta | Val | MG660667 | C16148T |
| 151 | D4j8 | Ud. Buryat | Kushun | MG660629 | - |
| 152 | D4p | Todzhi | Iiy | MG660705 | - |
| 153 | D4p | Todzhi | Adir-Kezhig | MG660710 | - |
| 154 | D5a2a1-16172! | Todzhi | Adir-Kezhig | MG660711 | T152C!, A16183C |
| 155 | F1b1b | Ket | Krasnoselkup | MG778671 | T10227C, C16179T, A16183C |
| 156 | F1b1b | Todzhi | Toora-Hem | MG660685 | A16183C, C16286T |
| 157 | F1b1b | Todzhi | Iiy | MG660689 | A16183C, C16286T |
| 158 | F1b1b | Todzhi | Iiy | MG660693 | G203A, C3204T, T10227C, C16179T, A16183C |
| 159 | F1b1b | Todzhi | Iiy | MG660700 | G203A, C3204T, T10227C, C16179T, A16183C |
| 160 | F1b1b | Todzhi | Adir-Kezhig | MG660708 | G203A, C3204T, T10227C, C16179T, A16183C |
| 161 | F1b1b | Todzhi | Adir-Kezhig | MG660713 | G203A, C3204T, T10227C, C16179T, A16183C |
| 162 | F1b1b | Todzhi | Adir-Kezhig | MG660714 | G203A, C3204T, T10227C, C16179T, A16183C |
| 163 | F1b1b | Todzhi | Adir-Kezhig | MG660717 | G203A, C3204T, T10227C, C16179T, A16183C |
| 164 | F1b1b | Tofalar | Alygdzher | MG660728 | A16183C, C16286T |
| 165 | F1b1e | Ud. Buryat | Alygdzher | MG660736 | A515G, A16183C |
| 166 | G1a1 | Todzhi | Toora-Hem | MG660682 | G4113A, A11908G, A14569G!, C15745A, C16260T |
| 167 | G2b2 | Todzhi | Adir-Kezhig | MG660715 | T152C!, T1709A, T5641C, A6647T, T11984C, A14569G!, T16093C |
| 168 | H101* | Ket | Maduika | MG778680 | T152C!, T9230C, A12290G, A15401C |
| 169 | H1ae3* | Ket | Kellog | MG778661 | T1189C, G8723A |
| 170 | H1b2 | Ket | V.Baikha | MG778677 | T6293C, G16129A!, A16183C |
| 171 | J1c2m | Ket | Sov. Rechka-Pakulikha | MG778682 | A3918G |
| 172 | M8a1 | Udegey | Bogorodskoye | MG660741 | A234G, A3744G |
| 173 | M8a1 | Udegey | Agzu/Ternei | MH807364 | A234G, A3744G |
| 174 | M8a2b | Nanai | Bogorodskoye | MG660742 | G12192A, G15106A |
| 175 | M8a3 | Ud. Buryat | Kushun | MG660627 | T146C!, T152C!, G5237A, T13488C, A16293G |
| 176 | M8a3 | Tofalar | Nerkha | MG660646 | T146C!, T152C!, G5237A, T13488C, A16293G |
| 177 | M8a3 | Tofalar | Nerkha | MG660650 | T146C!, T152C!, G5237A, T13488C, A16293G |
| 178 | M8a3 | Tofalar | V. Gutara | MG660745 | T146C!, T152C!, G5237A, T13488C, A16293G |
| 179 | M8a3 | Tofalar | V. Gutara | MG660747 | T146C!, T152C!, G5237A, T13488C, A16293G |
| 180 | M8a3 | Tofalar | V. Gutara | MG660749 | T146C!, T152C!, G5237A, T13488C, A16293G |
| 181 | M8a3 | Tofalar | V. Gutara | MG660750 | T146C!, T152C!, G5237A, T13488C, A16293G |

| | | | | | |
|---|---|---|---|---|---|
| 182 | M8a3a | Russian | Turukhansk | MG778679 | T146C!, T11260C, G13590A, A15684G, T16189C!, T16311C! |
| 183 | N2a | Ket | Kellog | MG778664 | T152C!, T1633C, T11722C, G12192A |
| 184 | U3b | Todzhi | Toora-Hem | MG660684 | G2652A, T13768C |
| 185 | U3b | Todzhi | Adir-Kezhig | MG660712 | G2652A, T10885C, T13768C |
| 186 | U4a1 | Ket | Kellog | MG778660 | T72C, A12950G, T14635C |
| 187 | U4a1 | Ket | Maduika | MG778665 | T72C, A12950G, T14635C |
| 188 | U4a1 | Ket | Farkovo | MG778655 | T72C, A12950G, T14635C |
| 189 | U4a1 | Ket | Kellog | MG778667 | T72C, A12950G, T14635C |
| 190 | U4a1 | Ket | Kellog | MG778657 | T72C, A12950G, T14635C |
| 191 | U4a1 | Ket | Pakulikha | MG778678 | T72C, A12950G, T14635C |
| 192 | U4a1 | Ket | Farkovo | MG778681 | T72C, A12950G, T14635C |
| 193 | U4a1 | Ket | Surgutikha | MG778658 | T72C, A12950G, T14635C |
| 194 | U4d2 | Ket | Surgutikha | MG778674 | C16111T, G16390A |
| 195 | U4d2 | Ket | Surgutikha | MG778675 | C16111T, G16390A |
| 196 | U4d2 | Ket | Vorogovo | MG778676 | A8596G, A12234G, C16189T!! |
| 197 | U4d2 | Mansi | Shaim | MG660638 | G10320A |
| 198 | U4d2 | Mansi | Urai | MG660642 | G10320A |
| 199 | U5a1d2 | Ket | Kellog | MG778669 | T146C!, T4823C |
| 200 | U5a1d2 | Ket | Baikha | MG660620 | T146C!, T4823C |
| 201 | U5a1d2 | Ket | Kellog | MG660621 | T146C!, T4823C |
| 202 | U5a1h* | Mansi | Aneyeva | MG660643 | C150T, C3192T, G3591A, A6895G, G12618A, A13966G, G16129A!, T16192C!, C16239T |
| 203 | U5a2a1b | Mansi | Verkhoturye | MG660639 | A9115G |
| 204 | U5a2b | Mansi | Shaim | MG660640 | C6713T, T16311C! |
| 205 | U5b2b | Ud. Buryat | Kushun | MG660623 | T8227C, T12696C, T15900C(hp), T16093C, C16223T!, C16234T(hp) |
| 206 | U5b2b | Tuvan | Bai-Tal | MG660668 | T7964C, T8077C, T9185G, G13968T, G14040A, T16189C!, C16260T |
| 207 | U5b2b | Tuvan | Bai-Tal | MG660669 | T7964C, T8077C, T9185G, G13968T, G14040A, A16183C, T16189C!, C16260T |
| 208 | Y1a1 | Ket | Yelogui | MG778663 | T3394C |
| 209 | Y1a1 | Ket | Yelogui | MG778656 | T3394C |
| 210 | Y1a1 | Ket | Kellog | MG778672 | T3394C |
| 211 | Y1a1 | Ket | Kellog | MG778673 | - |
| 212 | Z1 | Tofalar | Alygdzher | MG660740 | G207A, G251A, A3434G, T6425C, G9948A, T14180C, C16360T |
| 213 | Z1a | Koryak | Karaginskiy district | MK217935 | - |
| 214 | Z1a | Koryak | Karaginskiy district | MK217945 | - |
| 215 | Z1a1a | Ket | Kangatovo | MG778666 | - |
| 216 | Z1a2a | Koryak | Karaginskiy district | MK217916 | - |
| 217 | Z1a2a | Koryak | Karaginskiy district | MK217937 | - |
| 218 | Z1a2a | Koryak | Karaginskiy district | MK217940 | - |

**Supplementary Table 2.** List of ancient mtDNA samples used in this study. In bold are newly generated.

| # | Sample ID | mtDNA haplotype | Date | Culture/Arch.Period | Location/Cite | Country | Latitude | Longitude | Publication |
|---|---|---|---|---|---|---|---|---|---|
| 1 | DA344 | A+152+16362 | 4885 BP | LN | Ust'-Ida | Russia | 53,188889 | 103,368056 | Damgaard et al. 2018 |
| 2 | DA357 | A+152+16362 | 6713 BP | EN | Lokomotiv, Cis-Baikal | Russia | 52,286944 | 104,249167 | Damgaard et al. 2018 |
| 3 | RISE674 | A+152+16362 | 4061 BP (2281-1976 BC cal) | Okunevo, EMBA | Verkhni Askiz | Russia | 53,156486 | 90,207811 | Damgaard et al. 2018 |
| 4 | RISE680 | A+152+16362 | 4329 BP | Okunevo, EMBA | Uybat V | Russia | 53,708561 | 90,359808 | Damgaard et al. 2018 |
| 5 | RISE497 | A+152+16362 | 1400-900 BCE | Karasuk | Arban 1 | Russia | 52.954 | 90.187 | Allentoft et.al. 2015 |
| 6 | irk040 | A+152+16362 | N/A | Neolithic Cis-Baikal | Gorodische N 1 (Angara river, Irkutsk Oblast) | Russia | 53.220694 | 103.39475 | Kilinç et al. 2018 |
| 7 | irk025 | A+152+16362 | cal BC 2475 to 2335, cal BC 2325 to 2300 | Bronze Age Cis-Baikal | Sukhaja Pad' Buret' site, burial 2 | Russia | 52.983756 | 103.520719 | Kilinç et al. 2018 |
| 8 | brn001 | A+152+16362 | cal BC 5490 to 5465, cal BC 5400 to 5385 | Mesolithic Trans-Baikal | Izvestkovaja-1 site, burial 2 (Kuenga river, Sretensky District) | Russia | 52.230833 | 116.993333 | Kilinç et al. 2018 |
| 9 | irk030 | A10 | N/A | Neolithic Cis-Baikal | Korkino, burial 1 (Upper Lena river) | Russia | 54.373847 | 105.209969 | Kilinç et al. 2018 |
| 10 | N4a1 | A12a | cal BC 2830 to 2820, cal BC 2625 to 2475 | Late Neolithic Yakutia | Kyordyughen 2 (Central Yakutia) | Russia | 62.066667 | 132.333611 | Kilinç et al. 2018 |
| 11 | N4b2 | A12a | cal BC 2475 to 2295 | Late Neolithic Yakutia | Kyordyughen 1, burial 1 (Central Yakutia) | Russia | 62.066667 | 132.333333 | Kilinç et al. 2018 |
| 12 | I0562 | A16 | 400-300 BCE | Kazakhstan_Berel_IA | Berel | Kazakhstan | 49.3356 | 86.350367 | Unterlander et al. 2017 |
| 13 | RISE664 | A8a1 | 4272 BP (2459-2206 BC cal) | Okunevo, EMBA | Okunev Ulus | Russia | 53,547806 | 91,02565 | Damgaard et al. 2018 |
| 14 | RISE677 | A8a1 | 4419 BP (2831-2233 BC cal) | Okunevo, EMBA | Uybat III | Russia | 53,708561 | 90,359808 | Damgaard et al. 2018 |
| 15 | RISE681 | A8a1 | 4329 BP | Okunevo, EMBA | Uybat V | Russia | 53,708561 | 90,359808 | Damgaard et al. 2018 |
| 16 | RISE515 | A8a2 | 4197 BP (2340-2145 BC cal) | Okunevo, EMBA | Verkhni Askiz | Russia | 53,156486 | 90,207811 | Damgaard et al. 2018 |
| 17 | RISE667 | A8a2 | 4078.5 BP | Okunevo, EMBA | Verkhni Askiz | Russia | 53,156486 | 90,207811 | Damgaard et al. 2018 |
| 18 | RISE670 | A8a2 | 4088 BP (2141-1885 BC cal) | Okunevo, EMBA | Verkhni Askiz | Russia | 53,156486 | 90,207811 | Damgaard et al. 2018 |
| 19 | RISE673 | A8a2 | 4078.5 BP | Okunevo, EMBA | Verkhni Askiz | Russia | 53,156486 | 90,207811 | Damgaard et al. 2018 |
| 20 | RISE515 | A8a2 | 3810 BP (2340-2145 BC) | Okunevo | Verkhni Askiz | Russia | 53.153 | 90.194 | Allentoft et.al. 2015 |
| 21 | mak026 | C4 | cal BC 2895 to 2860, cal BC 2805 to 2755, cal BC 2720 to 2705 | Bronze Age Cis-Baikal | Makrushynskij burial site, burial 26 (Upper Lena river) | Russia | 53.876672 | 106.267394 | Kilinç et al. 2018 |
| 22 | irk057 | C4 | cal BC 2550 to 2535, cal BC 2490 to 2395, cal BC 2385 to 2345 | Bronze Age Cis-Baikal | Podostrozhnoe N 3 (Angara river, Irkutsk Oblast) | Russia | 53.220694 | 103.39475 | Kilinç et al. 2018 |
| 23 | **I1000** | **C4** | **2871-2497 calBCE (4100±40 BP, Poz-83436)** | **Glazkovskaya** | **Obkhoy, Kachugskiy district, Irkutsk region** | **Russia** | **54.0186686** | **105.4747222** | |
| 24 | DA247 | C4 | 6856 BP | EN | Shamanka II | Russia | 51,698333 | 103,703056 | Damgaard et al. 2018 |
| 25 | DA248 | C4 | 6815 BP | EN | Shamanka II | Russia | 51,698333 | 103,703056 | Damgaard et al. 2018 |
| 26 | DA249 | C4 | 7005 BP | EN | Shamanka II | Russia | 51,698333 | 103,703056 | Damgaard et al. 2018 |
| 27 | **I2072** | **C4+152** | **3959-3715 calBCE (5050±40 BP, Poz-83497)** | **Solentsy5_N** | **Solontcy 5, Foothhills of the Altai** | **Russia** | **52.4833** | **86.2167** | |
| 28 | RISE602 | C4+152 | 9/700 BC - AD 500/1000 | Iron Age | Sary-Bel | Russia | 50.615 | 84.459 | Allentoft et.al. 2015 |
| 29 | I0563 | C4a1a | 400-300 BCE | Kazakhstan_Berel_IA | Firsovo-XI, Ob bank, Altai province | Kazakhstan | 49.3356 | 86.350367 | Unterlander et al. 2017 |
| 30 | DA337 | C4a1a3 | 3871 BP | EBA | Shamanka II | Russia | 51,698333 | 103,703056 | Damgaard et al. 2018 |
| 31 | DA356 | C4a1a3 | 3854 BP | EBA | Ust'-Ida | Russia | 53,188889 | 103,368056 | Damgaard et al. 2018 |
| 32 | DA361 | C4a1a3 | 3854 BP | EBA | Ust'-Ida | Russia | 53,188889 | 103,368056 | Damgaard et al. 2018 |
| 33 | irk033 | C4a1a3 | cal BC 2920 to 2880 | Bronze Age Cis-Baikal | Chastaja Padi (Angara river, Irkutsk Oblast) | Russia | 52.989242 | 103.450869 | Kilinç et al. 2018 |
| 34 | irk008 | C4a1a3 | cal BC 5615 to 5485 | Mesolithic Trans-Baikal | Izvestkovaja-1 site, burial 1 (Kuenga river, Sretensky District) | Russia | 54.002744 | 105.710294 | Kilinç et al. 2018 |
| 35 | irk076 | C4a2a1 | cal BC 2275 to 2250, cal BC 2225 to 2220, cal BC 2210 to 2120 and cal BC 2090 to 2040 | Bronze Age Cis-Baikal | Shamanka 2, burial 3 (South Baikal) | Russia | 51.694478 | 103.70475 | Kilinç et al. 2018 |
| 36 | DA334 | C4a2a1 | 3764 BP | EBA | Shamanka II | Russia | 51,698333 | 103,703056 | Damgaard et al. 2018 |
| 37 | DA336 | C4a2a1 | 3817.5 BP | EBA | Shamanka II | Russia | 51,698333 | 103,703056 | Damgaard et al. 2018 |
| 38 | DA338 | C4a2a1 | 3817.5 BP | EBA | Shamanka II | Russia | 51,698333 | 103,703056 | Damgaard et al. 2018 |
| 39 | yak021 | C4b+16311! | cal BC 1385 to 1340, cal BC 1315 to 1195, cal BC 1140 to 1130 | Late Neolithic Yakutia | Pomazkino site, burial 2 (Kolyma river) | Russia | 67.916667 | 156.5 | Kilinç et al. 2018 |
| 40 | kra011 | C4b1 | cal BC 2295 to 2140 | Neolithic -Bronze? Krasnoyarsk Krai | Nefteprovod-2 site, burial 1 (Krasnoyarsk Krai) | Russia | 56.194736 | 95.819539 | Kilinç et al. 2018 |
| 41 | yak022 | C4b1 | cal BC 1940 to 1765 | Late Neolithic Yakutia | Kamenka 2 burial, individual 1 (Kolyma river) | Russia | 66.8 | 152.633333 | Kilinç et al. 2018 |
| 42 | yak023 | C4b1 | cal BC 1880 to 1690 | Late Neolithic Yakutia | Kamenka 2 burial, individual 2 (Kolyma river) | Russia | 66.8 | 152.633333 | Kilinç et al. 2018 |
| 43 | yak024 | C4b1 | in the same burial with yak022, yak023 | Late Neolithic Yakutia | Kamenka 2 burial, individual 3 (Kolyma river) | Russia | 66.8 | 152.633333 | Kilinç et al. 2018 |
| 44 | N3a | C4b3 | cal BC 790 to 730, cal BC 690 to 660, cal BC 650 to 540 | Iron Age Yakutia | Dyupsya burial (Central Yakutia) | Russia | 63.025 | 130.73 | Kilinç et al. 2018 |
| 45 | irk078 | C5 | cal BC 1260 to 1050 | Bronze Age Trans-Baikal | Okoshki 1, burial 23 (Zabaykalsky Krai) | Russia | 50.317222 | 118.275361 | Kilinç et al. 2018 |
| 46 | irk00x | C5+16093 | cal BC 6500 to 6435 | Mesolithic Trans-Baikal | Dzhylinda site (Chitinsuyu area) | Russia | 55.668922 | 115.872081 | Kilinç et al. 2018 |
| 47 | RISE684 | C5c | 4239 BP (2464-2141 BC cal) | EBA | Uybat V | Russia | 53,708561 | 90,359808 | Damgaard et al. 2018 |
| 48 | RISE685 | C5c | 4329 BP | EBA | Uybat V | Russia | 53,708561 | 90,359808 | Damgaard et al. 2018 |
| 49 | RISE718 | C5c | 4436 BP (2573-2348 BC cal) | EBA | Syda 5, Tumen | Russia | 54,371767 | 91,506856 | Damgaard et al. 2018 |
| 50 | RISE719 | C5c | 4450 BP | EBA | Syda 5, Tumen | Russia | 54,371767 | 91,506856 | Damgaard et al. 2018 |
| 51 | irk022 | D4b1c | cal BC 2455 to 2200 | Bronze Age Cis-Baikal | Ust'-Dolgoe site, burial 3 | Russia | 52.994322 | 103.420928 | Kilinç et al. 2018 |
| 52 | N5a | D4b1c | cal BC 4340 to 4235 | Middle Neolithic Yakutia | Onnyos burial (Amga river) | Russia | 60.458333 | 131.091667 | Kilinç et al. 2018 |
| 53 | DA358 | F1b | 4169 BP | EBA | Kurma XI | Russia | 53,179167 | 106,962778 | Damgaard et al. 2018 |
| 54 | DA360 | F1b | 4158 BP | EBA | Kurma XI | Russia | 53,179167 | 106,962778 | Damgaard et al. 2018 |
| 55 | DA253 | F1b1+152 | 6329 BP | EN | Shamanka II | Russia | 51,698333 | 103,703056 | Damgaard et al. 2018 |
| 56 | DA335 | F1b1b | 3818 BP | EBA | Shamanka II | Russia | 51,698333 | 103,703056 | Damgaard et al. 2018 |
| 57 | irk036 | F1b1b(2) | cal BC 2885 to 2835, cal BC 2815 to 2665 | Bronze Age Cis-Baikal | Glazkovo (Angara river, Irkutsk Oblast) | Russia | 52.286378 | 104.260106 | Kilinç et al. 2018 |
| 58 | RISE553 | F1b1b2 | 2731 BP (926-815 BC) | LBA | Afontova Gora, Krasnoyarsk | Russia | 56.016 | 92.866 | Allentoft et.al. 2015 |
| 59 | irk068 | F1b1b3 | N/A | Neolithic Cis-Baikal | Shishkino N 1 (Upper Lena river) | Russia | 54.006611 | 105.681531 | Kilinç et al. 2018 |
| 60 | RISE554 | F1b1b3 | 2782 BP (1005-844 BC) | LBA | Afontova Gora, Krasnoyarsk | Russia | 56.016 | 92.866 | Allentoft et.al. 2015 |
| 61 | I0211 | U4a | 6773-5886 BCE | Karelia_HG | Yuzhnyy Oleni Ostrov, Karelia | Russia | 61.65 | 35.65 | Mathieson et.al. 2015 |
| 62 | **I0992** | **U4a(3)** | **5002-4730 calBCE (5990±50 BP, Poz-83428)** | **Novosibirsk_N** | **Korchugan-1, Novosibirsk region** | **Russia** | **56.4666647** | **76.3** | |
| 63 | **I2074** | **U4a(3)** | **5602-5376 calBCE (6520±40 BP, Poz-83514)** | **Kemerovo_N** | **Vas'kovo-4, burial 1, Intermountain basin between spurs of Altai and Sayan Mountains** | **Russia** | **55.0537** | **85.0966** | |
| 64 | I0231 | U4a1 | 2921-2762 calBCE (4260±30 BP, Beta-392487) | Yamnaya_Samara | Ekaterinovka, Southern Steppe, Samara | Russia | 52.42 | 48.24 | Mathieson et.al. 2015 |
| 65 | SBj | U4a1 | 8963-8579 calBP | Mesoltihic Scandinavian Hunter-gatherer | Stora Bjers | Sweden | 57.8167 | 18.53 | Gunther et.al. 2018 |
| 66 | SF12 | U4a1 | 9033–8757 calBP | Mesoltihic Scandinavian Hunter-gatherer | Stora Förvar | Sweden | 57.2853 | 17.9706 | Gunther et.al. 2018 |
| 67 | I0434 | U4d | 5200-4000 BCE | Samara_Eneolithic | Khvalynsk II, Volga River, Samara | Russia | 52.22 | 48.1 | Mathieson et.al. 2015 |
| 68 | RISE500 | U4d1 | 1700-1500 BC | Andronovo | Kytmanovo | Russia | 53.456 | 85.447 | Allentoft et.al. 2015 |
| 69 | I0124 | U5a1d | 5657-5541 calBCE (6680±30 BP, Beta-392490) | Samara_HG | Lebyazhinka IV, Sok River, Samara | Russia | 53.4 | 50.4 | Mathieson et.al. 2015 |
| 70 | Hum2 | U5a1d | 9452-9275 calBP | Mesoltihic Scandinavian Hunter-gatherer | Hummervikholmen, Søgne archipelago, Southern Norway | Norway | 58.064 | 7.7438 | Gunther et.al. 2018 |
| 71 | Steigen | U5a1d | 5950-5764 calBP | Mesoltihic Scandinavian Hunter-gatherer | Måløya, Steigen, Northern Norway | Norway | 67.81 | 14.6818 | Gunther et.al. 2018 |
| 72 | RISE502 | U5a1d | 3140 BP (1496-1306 BC) | Karasuk | Bystrovka | Russia | 51.909 | 88.574 | Allentoft et.al. 2015 |
| 73 | RISE240 | U5a1d1 | 4160 BP (2880-2632 BC) | Yamnaya | Sukhaya Termista I | Russia | 46.58 | 43.678 | Allentoft et.al. 2015 |
| 74 | **I2068** | **U5a1d2** | **420-565 calCE (1560±30 BP, Poz-83507)** | **Siberia_IA** | **Tepsei III, kurgan 2, Minusinskaya intermauntain basin, Sayan Mountain** | **Russia** | **53.9634** | **91.5587** | |
| 75 | RISE546 | U5a1d2b | 3000-2400 BC | Yamnaya | Temrta IV | Russia | 46.539 | 43.699 | Mathieson et.al. 2015 |
| 76 | **I0991** | **Z1** | **5206-4805 calBCE (6060±50 BP, Poz-83427)** | **Novosibirsk_N** | **Korchugan-1, Novosibirsk region** | **Russia** | **56.4666647** | **76.3** | |
| 77 | **I0998** | **Z1a1** | **2835-2472 calBCE (4040±35 BP, Poz-83426)** | **Serovskaya** | **Khuzhir, Lake Baikal, Olkhon Island, Irkutsk region** | **Russia** | **53.193333** | **107.343889** | |
